# Combination Immunotherapy Enhances Serial Radiofrequency Ablation Induced Systemic Antitumor Immunity in Pancreatic Cancer

**DOI:** 10.64898/2026.04.05.713683

**Authors:** Lincoln N. Strickland, MacKenzie V. Demmel, Wendao Liu, Alyssa M. Waller, Shwetapadma Dash, Khadija Turabi, Urvinder Kaur Sardarni, Nicolette R. Mardik, Casey J. Van Kirk, Baylee O’Brien, Julie Rowe, Putao Cen, Kelsey A. Klute, Jesse L. Cox, Zhongming Zhao, Sunil R. Hingorani, Curtis J. Wray, Nirav C. Thosani, Jennifer M. Bailey-Lundberg

**Affiliations:** Department of Pathology, Microbiology, and Immunology, University of Nebraska Medical Center, Omaha, NE; Fred & Pamela Buffett Cancer Center, University of Nebraska Medical Center, Omaha, NE; Cancer Research Graduate Program, The Eppley Institute, Fred & Pamela Buffett Cancer Center, University of Nebraska Medical Center, Omaha, NE; The University of Texas MD Anderson Cancer Center UTHealth Houston Graduate School of Biomedical Sciences, Houston, Texas; Center for Precision Health, McWilliams School of Biomedical Informatics, The University of Texas Health Science Center at Houston, Houston, TX; Biochemistry and Molecular Biology Graduate Program, University of Nebraska Medical Center, Omaha, NE; Department of Internal Medicine, The University of Texas Health Science Center at Houston, Houston, TX; Department of Internal Medicine, University of Nebraska Medical Center, Omaha, NE; Pancreatic Cancer Center of Excellence, University of Nebraska Medical Center, Omaha, NE; Department of Surgery, The University of Texas Health Science Center at Houston, Houston, TX; Center for Interventional Gastroenterology, The University of Texas Health Science Center, Houston, TX

**Keywords:** Radiofrequency ablation, immunotherapy, therapeutic resistance, CSF1R

## Abstract

Thermal ablation is increasingly used for local control of pancreatic ductal adenocarcinoma (PDAC), but its capacity to induce systemic antitumor immunity and the mechanisms limiting this response remain incompletely defined. Using a bilateral *LSL-Kras^G12D/+;^ LSL-Trp53^R172H/+^*; *Pdx1-Cre (KPC)* flank tumor model, we show serial radiofrequency ablation (RFA) enhances local tumor control and induces a robust abscopal response. This effect was associated with increased activation of CD8⁺ T cells and natural killer cells, and was abrogated by CD8⁺ T cell depletion. Single-cell RNA sequencing revealed expansion of cytotoxic immune programs alongside induction of a CSF1R-driven myeloid response consistent with adaptive immune resistance. Although combinatorial blockade of PD-L1 and CD73 augmented systemic antitumor responses, and the addition of CSF1R inhibition in this context further enhanced both local and distant tumor control. These findings identify a CSF1R-dependent myeloid resistance program which constrains ablation-induced systemic immunity and demonstrate that rational combination immunotherapy can potentiate the systemic efficacy of tumor ablation in PDAC.

**STATEMENT OF TRANSLATIONAL RELEVANCE:** Radiofrequency ablation is a local thermal therapy that induces tumor destruction and modulation of the tumor microenvironment used for treatment of many solid tumors, including pancreatic cancer. Serial thermal ablations induce a CD8+ T cells and NK cell antitumor immunological response; however, *Nt5e* (CD73) is increased in tumor cells, PD-1 is increased in CD8+ T cells, and serial ablations increase a CSF1R+ macrophage cluster in ablated tumors. Combinational targeting of CD73, PD-L1, and CSF1R alleviates CSF1R-dependent therapeutic resistance and enhances local and systemic tumor control. This study identifies targetable countervailing mechanisms to local ablation therapy and provides a rationale for combinatorial strategies integrating serial tumor ablation with systemic immunomodulatory agents. Future prospective studies should evaluate the influence of timing, dosage, and sequence of ablation on immune priming to improve clinical interventions in pancreatic cancer.

## INTRODUCTION

Radiofrequency ablation (RFA) is an established locoregional therapy that induces tumor destruction through thermal coagulative necrosis and is widely used for the treatment of solid malignancies, including hepatic, renal, and pancreatic tumors^1–9^. In the treatment of pancreatic tumors, endoscopic ultrasound-guided radiofrequency ablation (EUS-RFA) has emerged as a minimally invasive approach delivering targeted thermal energy to pancreatic lesions while minimizing injury to surrounding structures. Early clinical studies demonstrate feasibility and safety of EUS-RFA for pancreatic tumors, including pancreatic ductal adenocarcinoma (PDAC) and pancreatic neuroendocrine tumors, suggesting this strategy may provide local tumor control in patients who are not candidates for surgical resection^10–12^. Beyond its direct cytotoxic effects, accumulating evidence indicates thermal ablation can also modulate the tumor immune microenvironment (TIM)^13–15^.

RFA induced thermal injury leads to rapid tumor cell death and the release of tumor-associated antigens, danger-associated molecular patterns, and inflammatory cytokines that can promote immune activation. Preclinical studies have demonstrated RFA can enhance antigen presentation, stimulate dendritic cell maturation, and promote infiltration of cytotoxic lymphocytes within the tumor microenvironment (TME)^16,17^. In some settings, these immune responses extend beyond the site of ablation, generating systemic antitumor immunity and tumor regression at distant untreated sites resembling an abscopal effect^14,18,19^. However, the magnitude and durability of these immune responses are often limited, particularly in solid tumors such as PDAC that are characterized by profound immune suppression and resistance to immunotherapy^20,21^.

Importantly, most preclinical studies investigating the immunologic consequences of thermal ablation have focused on single-treatment paradigms, even though patients frequently undergo multiple ablation procedures due to incomplete tumor destruction, recurrence, or staged treatment strategies^22,23^. Serial tumor injury may amplify inflammatory signaling and antigen release, potentially enhancing immune priming and immune cell recruitment within the TME. Simultaneously, tissue injury can promote hypoxia, alter the stiffness and subtypes of cancer-associated fibroblasts (CAFs), and enhance the recruitment of immunoregulatory myeloid populations, including tolerogenic or antitumor neutrophils and macrophages, which shape the balance between antitumor immunity and immune suppression^14,24–26^. Among these pathways, the colony stimulating factor 1/colony stimulating factor 1 receptor (CSF1/CSF1R) signaling axis plays a central role in regulating macrophage recruitment, differentiation, and polarization within the TME and has been implicated in resistance to immunotherapy across multiple solid tumors, including PDAC^27–29^.

Given the highly immune suppressive nature of PDAC, strategies capable of converting the TME into an immunologically active state may be required to sensitize tumors to immunotherapy^30^. EUS-guided locoregional therapies such as RFA may provide such an opportunity by inducing immunogenic tumor cell death and promoting immune infiltration. However, the extent to which serial thermal ablation reshapes the tumor immune landscape and generates systemic antitumor immunity remains poorly understood. Moreover, how ablation-induced immune remodeling interacts with key immune suppressive pathways, including macrophage signaling and immune checkpoint regulation, has not been fully defined.

In this study, we investigated how serial thermal ablation using RFA influences tumor progression and the TIM using a bilateral flank PDAC tumor model that enables simultaneous evaluation of local and systemic immune responses. Through histologic, single-cell transcriptomic, and immunologic analyses, we examined how sequential ablation events remodel immune cell populations and inflammatory signaling networks within the RFA-treated and contralateral tumors. We show serial thermal ablation using RFA in combination with targeting PD-L1, CD73 and CSF1R significantly restrains the growth of treated and contralateral tumors. These data are encouraging given the challenges of generating durable responses to immunotherapy in solid tumors including PDAC.

## METHODS

### Preclinical KPC Subcutaneous Tumor Model

All mouse model procedures followed UNMC’s IACUC protocol and adhere to ARRIVE guidelines. 500,000 KPC cells were prepared in a PBS:Matrigel mix (1:1) and injected subcutaneously in both flanks of 8-week-old male C57BL/6 mice. Sex as a biological variable: male mice were used for these experiments as the KPC line was derived from male mice and tumor growth rates are more variable in female mice. Tumor size was measured and recorded twice per week with a vernier caliper. Tumor volume was calculated as ((length × width × width)/2) in mm^3^. KPC cells were derived in the Tuveson Lab from *Kras^LSL-G12D/+^;Trp53^LSL-R172H/+^;Pdx1-Cre* mice, which develop PDAC. Statistical analysis was performed using a one-way ANOVA or student’s *t* test in Prism GraphPad software.

### Radiofrequency Ablation

RFA was performed when the tumors were between 200-500 mm^3^ 14, 18, and 21 days after implantation. Mice were anesthetized with isoflurane via a nose cone. Ablation was performed on the larger tumor. Skin around the ablation site was cleaned with 3 alternating rounds of 70% ethanol and iodine and a 2.5 mg/kg dose of bupivacaine (NDC 0409-1162-18, Hospira Inc.) and 5 mg/kg dose of meloxicam (NDC 13965-559-20, Vet One) was administered through subcutaneous injection prior to ablation. A small incision was made in skin at the center of the tumor and the Habib EUS-RFA probe was inserted. Ablation was performed for 10 seconds and the average power delivered was 0.5 W (watts). Mice were observed for signs of pain post-ablation. The sham control mice were transferred to the procedure room, placed under anesthesia, received bupivacaine and meloxicam, an incision was made, the probe was inserted, but no ablation was performed.

### Hematoxylin and Eosin Staining and Quantification

Hematoxylin and eosin (H&E) staining was performed as previously described^5^. Statistical analysis was performed using a student’s *t* test in Prism GraphPad software.

### Immunohistochemistry and Quantification

Immunohistochemistry (IHC) staining and quantification was performed as previously described^5^. Statistical analysis was performed using a one-way ANOVA in Prism GraphPad software. Antibodies and dilutions used are in **Supplemental Table 1**.

### Immunofluorescence

Slides were baked at 60°C for 30 minutes, deparaffinized in histoclear for 9 minutes, rehydration was performed by 2-minute washes in graded alcohols at 100%, 95%, then 70%. Slides were placed in 1x PBS for 6 minutes, in PBST for 15 minutes, and 1x PBS for 10 minutes. Antigen retrieval was performed based on antibody recommendations. Slides cooled completely before 10 minutes in 1x PBS. Slides were blocked in 10% FBS in 1x PBST for 60 minutes before primary antibody incubation overnight in a humidified chamber at 4°C. Slides were washed 3 times with PBST for 10 minutes each. Secondary antibody was diluted in blocking solution and incubated for 90 minutes, protected from light. Quenching (103710-194, Novus Biologicals) was performed for 5 minutes. Hoechst staining (MSPP-62249, Invitrogen) was performed for 5 minutes before slides were mounted with VECTASHIELD vibrance antifade mounting media (H-1000-10, Vector Laboratories) and imaged. Antibodies and dilutions used are in **Supplemental Table 1**. Statistical analysis was performed using a one-way ANOVA in Prism GraphPad software.

### Single cell RNA-seq and Analysis

For the one RFA group, tumor tissue from two mice was pooled into a single sample. For the three RFA group, tumor tissue from four mice was pooled into two samples (two mice per sample). Consequently, a total of three processed samples were subjected to paired scRNA-seq and scTCR-seq. Processing was performed as previously described^14^ and submitted to the Cancer Genomics Center at UTHealth. Raw data was processed from paired scRNA library and single-cell TCR library Cell Ranger (v7.0.0) multi pipeline. RNA library was aligned to mm10-2020-A reference genome, and TCR libraries were aligned to GRCm38-alts-ensembl-7.0.0 provided by 10× Genomics. Due to few T cells, TCR output was not included in the analysis. Subsequently, detailed QC metrics were computed and assessed using Seurat package (v5.4.0)^31^. Cells with less than 200 genes, more than 7,000 genes, or more than 20% mitochondrial gene counts were filtered out to ensure data quality. Raw counts were normalized with the function *NormalizeData*. Doublet detection was performed using scDblFinder (v1.24.10)^32^. Across the three samples, doublet rates were low (mean 4.5%, range 3.8–5.6%, corresponding to approximately 45 cells per sample). Singlet-filtered and unfiltered datasets were compared and found to produce concordant cluster assignments, as shown in **Supplemental Fig. S1**. Thus, given the limited cell numbers per sample (700–1,300 cells) and the low doublet burden, all cells were retained for downstream analysis with doublet classification preserved as metadata. After QC filtering, there were 728 cells remaining in the one RFA group, 1306 cells in the first three RFA sample, and 1186 in the second three RFA sample. Data were scaled using the *ScaleData* function. Principal component analysis (PCA) was then performed on the 2,000 highly variable genes. Samples were integrated using harmony (v2.0.5)^33^. The *FindNeighbors* function was used with the first 35 PCs to build the nearest neighbor graph and the *FindClusters* function was used with a 0.6 resolution to identify cell clusters. The UMAP method was employed for dimensionality reduction and 2D visualization of cell clusters. Differential expression analysis was performed using the *FindMarkers* function with Wilcoxon ranked-sum test. Cell types were annotated by examining the expression of canonical cell markers in cell clusters. The difference in gene expression between two conditions of a cell type was tested using Wilcoxon ranked-sum test and adjusted using the Benjamini-Hochberg procedure. Heatmaps were generated using the package *ComplexHeatmap* (v2.26.1)^34^. The package ggplot2 (v4.0.3)^35^ was used to generate violin plots for these gene expression levels. The difference in cell proportion between two conditions of a cell type was tested using Fisher exact test. P-values were adjusted using the Benjamini-Hochberg procedure.

### Cytokine Array Proteome Profiler

The cytokine array proteome profiler was performed as previously described^14^. Statistical analysis was performed using a student’s t-test in Prism GraphPad software.

### PLX3397 Administration

PLX3397 (HY-16749, MedChem Express) was dissolved in 10% DMSO (D8418, Sigma-Aldrich) and 90% 20% SBE-β-CD (HY-17031/CS-0731, MedChem Express) in saline. PLX3397 was administered through oral gavage at a 50 mg/kg dose daily beginning 13 days after tumor implantation on the day before the first RFA treatment until the experiment endpoint. The vehicle control was 10% DMSO in 20% SBE-β-CD in saline.

### Anti-PD-L1 Administration

Anti-PD-L1 antibody (B7-H1, BioXCell) was diluted in InVivo Pure pH 7.0 dilution buffer (IP0070, BioXCell). Anti-PD-L1 was administered through intraperitoneal injections at a 200 μg dose beginning 14 days after tumor implantation, on the day of the first RFA treatment and was given every other day until the experiment endpoint. The anti-IgG2b isotype control (BE0090, BioXCell) was diluted in InVivo Pure pH 7.0 dilution buffer (IP0070, BioXCell) and administered at the same dose and schedule as anti-PD-L1.

### Quemliclustat Administration

Quemliclustat (HY-125286, MedChemExpress) was dissolved in 20% DMSO (D8418, Sigma-Aldrich) and 90% 20% SBE-β-CD (HY-17031/CS-0731, MedChem Express) in saline. Quemliclustat was administered through oral gavage at a 10 mg/kg dose beginning 14 days after tumor implantation, on the day of the first RFA treatment and was given every other day until the experiment endpoint. The vehicle control was 10% DMSO in 20% SBE-β-CD in saline.

### Anti-CD8α Administration

Anti-CD8α antibody (BP0061, BioXCell) was diluted in InVivo Pure pH 7.0 dilution buffer (IP0070, BioXCell). Anti-CD8α was administered through intraperitoneal injections at a 200 μg dose beginning 13 days after tumor implantation, on the day before the first RFA treatment and was given every other day until the experiment endpoint. The anti-IgG2b isotype control (BP0090, BioXCell) was diluted in InVivo Pure pH 7.0 dilution buffer (IP0070, BioXCell) and administered at the same dose and schedule as anti-CD8α.

### Flow Cytometry Validation of CD8α Depletion

Whole blood was collected at the time of euthanasia and centrifuged for 5 minutes at 500 g. Serum was collected. 1 mL of 1x PBS was added and samples were centrifuged for 5 minutes at 500 g. Remaining serum was collected. Samples were incubated in 1 mL RBC lysis buffer (420301, BioLegend) for 10 minutes, then 1 mL 1x PBS was added and samples were centrifuged for 5 minutes at 500 g. Samples were incubated in Live/Dead Fix Blue (Thermo Fisher Scientific, L34962) for 15 minutes then washed with 1 mL 1x PBS and centrifuged for 5 minutes at 500 g. Fc receptor block (BD Biosciences, 553142) was added for 5 minutes on ice, then anti-CD8α antibody (100714, BioLegend) was incubated for 15 minutes at room temperature. Samples were washed with 1x PBS and centrifuged for 5 minutes at 500 g. Samples were resuspended in .5% BSA/PBS. Data was acquired using a BD LSRFortessa and analyzed with FlowJo v10.10.0.

### Clinical Samples

This study represents an approved institutional review of a prospective longitudinal cohort and phase 2 clinical trial (PANCARDINAL-1) involving EUS (protocols HSC-MS-21-0066 and HSC-MS-18-0192). All procedures involving human participants were conducted in compliance with the ethical standards set by the UTHealth Houston Committee for the Protection of Human Subjects and adhered to the principles outlined in the 1961 Declaration of Helsinki. Eligible participants were required to have histologically confirmed PDAC. Based on physician’s choice, patients received chemotherapy consisting of modified FOLFIRINOX, gemcitabine and nab-paclitaxel with or without cisplatin. In a subset of patients, EUS-RFA was performed after one month of chemotherapy. Under EUS guidance, an electrode needle probe was advanced into the pancreatic ductal adenocarcinoma via either a transduodenal or transgastric approach. Prophylactic antibiotics and nonsteroidal anti-inflammatory drugs were not routinely administered. Color Doppler imaging was used to minimize the risk of vascular injury. Ablation was continued until electrical impedance reached 200 Ohms or demonstrated a rapid increase. Treatment effect was confirmed by real-time sonographic changes in tumor appearance, including alterations in echogenicity and evidence of tissue liquefaction. Patients typically continued with RFA sessions until the interventional gastroenterologist evaluated that pancreatic tumor could no longer be amenable to further RFA. Clinical, demographic, and perioperative data if patient underwent surgical resection of PDAC tumor were systematically collected. Radiologic response was assessed by comparing pre- and post-RFA axial computed tomography or magnetic resonance imaging using RECIST 1.1 criteria. Additionally, tumor characteristics were evaluated through changes in Hounsfield units before and after treatment. Size of the pancreatic tumor was also measured and recorded for each RFA session.

## RESULTS

### Serial RFA Remodels the Tumor Immune Microenvironment in Treated and Contralateral Tumors

While we have previously demonstrated significant reductions in the size of one RFA-treated tumors at day four compared to a sham treatment^14^ in a bilateral syngeneic KPC flank tumor model, we began here by utilizing this model to investigate the effect of serial thermal RFA sessions (**Fig. 1A**). We compared the tumor volumes of locally ablated tumors and distant contralateral tumors 25 days after tumor implantation of control untreated tumors, mice receiving 1 RFA session on day 14, and mice receiving either 3 serial sham treatments or 3 serial RFA treatments given 4 days apart beginning on day 14. Serial RFA every four days significantly reduced tumor the volume of the ablated tumors compared to one RFA session and to sham treated tumors (**Fig. 1B**). However, the contralateral tumors were significantly reduced compared to untreated control tumors, tumors receiving one ablation, and tumors receiving three serial sham treatments (**Fig. 1C**). This indicates that while serial RFA is locally beneficial, the abscopal effect is further enhanced by serial ablations.

**Figure 1.**
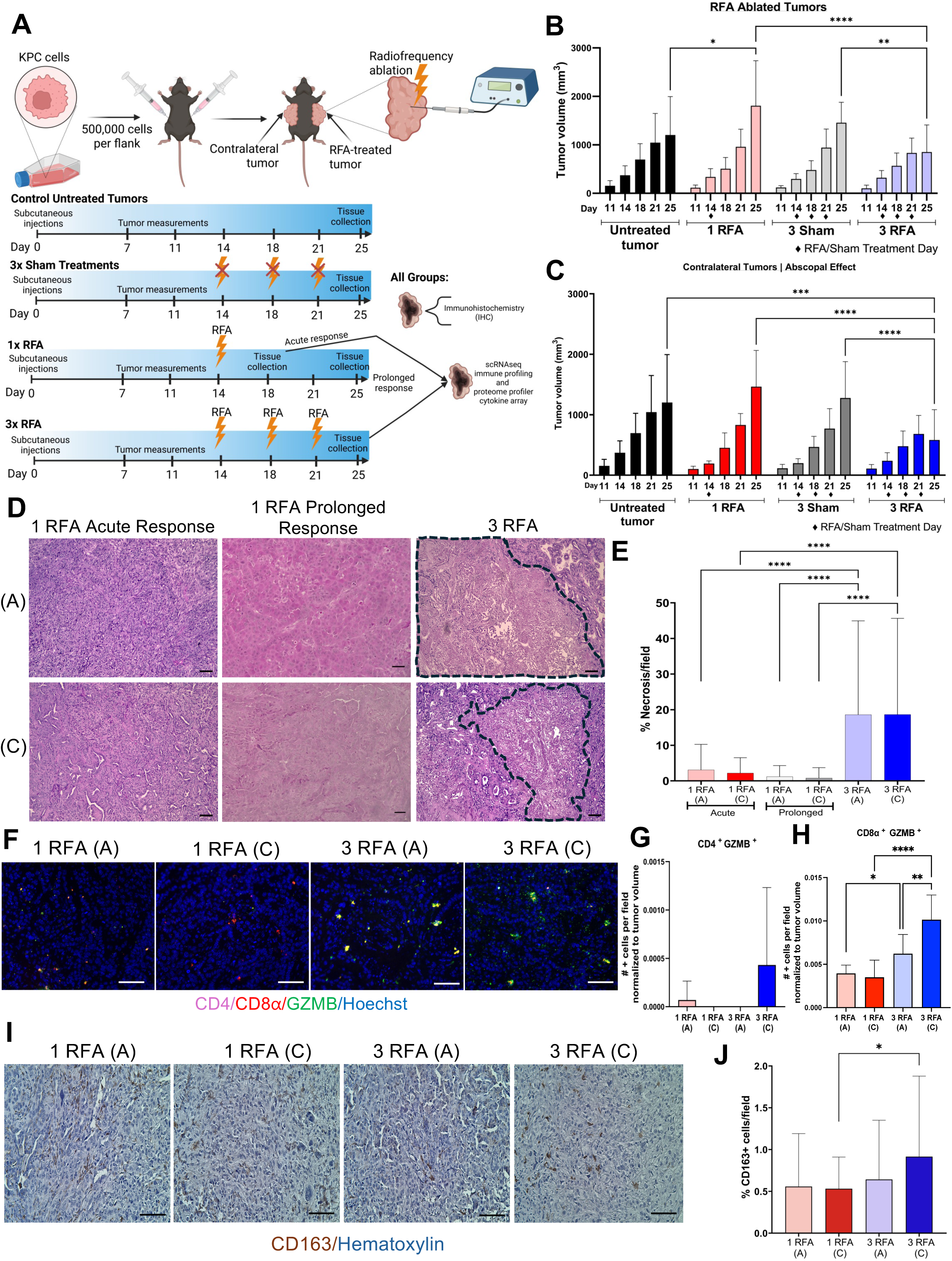
Serial radiofrequency ablation remodels the tumor immune microenvironment in treated and contralateral tumors. **A)** Experimental design created using BioRender. **B)** On the treated tumor, serial RFA sessions significantly reduce tumor volume at day 25 compared to control untreated tumors (*p=0.0163), 3 sham treatments (**p=0.0058), and to 1 RFA session (***p<0.0001). **C)** On the contralateral tumor, serial RFA sessions significantly reduce tumor volume at day 25 compared to control untreated tumors (***p=0.0001), 3 sham treatments (****p<0.0001), and to 1 RFA session (***p<0.0001). For **B-C**, n=18 for untreated tumors, n=19 for 1 RFA, n=23 for 3 RFA, and n=10 for 3 Sham. **D)** Representative 10x hematoxylin and eosin images, necrosis outlined in dashed line. **E)** Necrosis is significantly increased in 3 RFA ablated tumors compared to 1 RFA acute and prolonged ablated tumors (****p<0.0001). Necrosis is significantly increased in 3 RFA contralateral tumors compared to 1 RFA acute and prolonged contralateral tumors (****p<0.0001). **F)** Representative 20x immunofluorescence images of CD4 (purple), CD8α (red), Granzyme B (green), Hoechst nuclear stain (blue). **G)** CD4+GZM+ cells are not increased in 3 RFA contralateral tumors. **H)** CD8α+GZM+ cells are significantly increased in the three RFA contralateral tumors compared to 1 RFA contralateral (****p<0.0001) and 3 RFA RFA-treated tumors (**p=0.0046). n=8 fields analyzed per group. **I)** Representative 20x images of CD163 staining. **J)** CD163 staining is significantly increased in 3 RFA contralateral tumors compared to 1 RFA contralateral tumors (*p=0.0155). Scale bars 50 μm.

Histopathologic evaluation of tumors using hematoxylin and eosin (H&E) staining revealed substantially larger areas of necrosis (dashed black lines) in tumors receiving serial ablation compared to tumors treated with one RFA and examined both four days (acute response) and eleven days (prolonged response) after the ablation procedure (**Fig. 1D and E**). Notably, significantly increased necrosis was also observed in contralateral tumors from the three RFA treatment group compared with contralateral tumors from the one RFA treatment group examined at both the acute and prolonged timepoints indicating a more pronounced abscopal response after serial thermal ablation (**Fig. 1E**). Additionally, apoptosis (cleaved caspase 3 (CC3)) was significantly increased in both three RFA-treated and contralateral tumors compared to one RFA-treated and contralateral tumors (**Supplemental Fig. S2A and B**). Given the reduction of tumor volume and increase in tumor cell death in the ablated and contralateral tumors following serial RFA, we began to further characterize the tumors using IHC and multiplexed immunofluorescence (IF) staining to examine changes in the tumor immune microenvironment following serial ablation.

Recent preclinical studies demonstrate neutrophils to be a dominant innate immune cell infiltrating RFA-treated tumors^13,14^. To further characterize innate immune infiltration, IHC was performed for neutrophils using GR1 and myeloperoxidase (MPO) (**Supplemental Fig. S2A**). GR1^+^ cells were significantly increased in serial RFA-treated and contralateral tumors compared to one RFA tumors (**Supplemental Fig. S2C**). Interestingly, GR1 expression was significantly higher in three RFA contralateral tumors than in the corresponding serial RFA-treated tumors and MPO staining was also significantly increased in three RFA-treated contralateral tumors (**Supplemental Fig. S2D**).

To characterize cytotoxic lymphocyte activity, multiplex immunofluorescence (IF) staining was performed for CD4, CD8α, and GZMB (**Fig. 1F; Supplemental Fig. S3**). While not significant, quantitative analysis revealed the presence of CD4⁺GZMB⁺ cells in contralateral, but not RFA-treated tumors following serial RFA treatment (**Fig. 1G).** In contrast, CD8α⁺GZMB⁺ cytotoxic T cells were significantly enriched in both the ablated and contralateral tumors from the three RFA group compared to the respective tumors from the one RFA-treated group (**Fig. 1H**). Interestingly, the most CD8α⁺GZMB⁺ cytotoxic T cells were present in the serial RFA contralateral tumors. Collectively, these findings demonstrate serial thermal ablation using RFA promotes a more immunologically active TME in both local and distant tumors.

IHC was also performed to characterize infiltrating macrophage populations using CD68, inducible nitric oxide synthase (iNOS), and CD163 (**Fig. 1I, Supplemental Fig. S2A**). We observed a significant reduction in CD68^+^ cells in the three RFA-treated tumors compared to one RFA-treated and serial RFA contralateral tumors (**Supplemental Fig. S2E**). However, there was enhanced inflammatory iNOS+ macrophage infiltration following serial ablation in both the ablated and contralateral tumors compared to single RFA treated and contralateral tumors (**Supplemental Fig. S2F**). In addition, iNOS+ cells were significantly increased in three RFA contralateral tumors compared with the corresponding RFA-treated tumors (**Supplemental Fig. S2F**). Finally, CD163 expression, a marker associated with immune suppressive macrophages, was significantly increased in contralateral tumors from the three RFA treatment group compared to one RFA contralateral tumors (**Fig. S2I and J**). Together, these findings demonstrate serial RFA not only increases tumor necrosis at the treatment site but also significantly alters the immune landscape both locally and systemically.

To evaluate the functional role of CD8⁺ T cells, we performed a depletion experiment (**Supplemental Fig. S4A**). As assessed through flow cytometry, anti-CD8α treatment efficiently reduced systemic CD8⁺ T cell populations in whole blood **(Supplemental Fig. S4B and E).** Anti-CD8α T cell depletion significantly accelerated tumor growth in both treated and contralateral tumors following two RFA treatments but did not affect final tumor volumes after a third RFA **(Supplemental Fig. S4C and D)**, indicating CD8⁺ T cells mediate acute tumor control, but the effect is not as sustained after the third ablation.

### Single-Cell RNA Sequencing Reveals Serial RFA-Dependent Tumor Cell States and Enhanced Cytotoxic Immune Signatures

To define how serial thermal ablation reshapes the cellular composition of the tumor microenvironment, we performed single-cell RNA sequencing (scRNA-seq) on tumors following either one or three RFA treatments. Dimensionality reduction using uniform manifold approximation and projection (UMAP) revealed substantial cellular heterogeneity across the tumors (**Fig. 2A, Supplemental Fig. S5A**). In total, 14 transcriptionally distinct clusters were identified and annotated into 8 major cell types based on canonical marker gene expression (**Supplemental Fig. S5B)**. These included five tumor cell clusters (clusters 0, 3, 6, 8, and 9), two macrophage clusters, two neutrophil clusters, and single clusters corresponding to erythroid cells, endothelial cells, classical dendritic cells, T cell/natural killer (NK) cells, and B cells.

**Figure 2.**
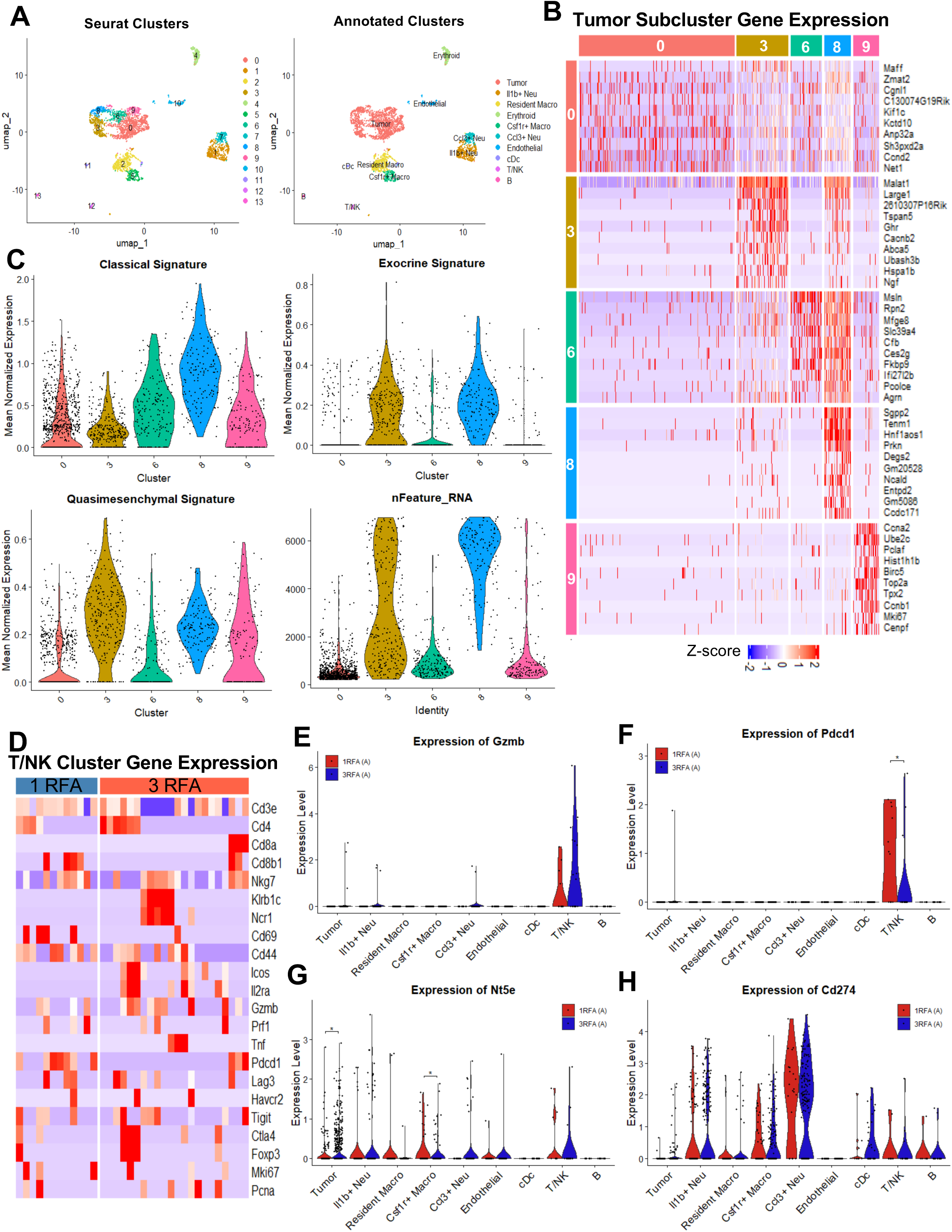
scRNA-seq reveals serial RFA-dependent tumor cell states and enhanced cytotoxic immune signatures. **A)** uMAPs of identified cell clusters and cell types from single-cell RNA-sequencing analysis. There are 14 cell clusters and 8 cell types (5 tumor clusters, 2 macrophage clusters, 2 neutrophil clusters, and an erythroid, endothelial, classical dendritic cell, T/natural killer cell, and B cell cluster). **B)** Heat map showing top differentially expressed genes in the 5 tumor clusters (0, 3, 6, 8, and 9). **C)** Violin plots showing tumor cluster gene expression related to a classical, exocrine, or quasimesenchymal signature and nFeature RNA plot. **D)** Heat map showing top differentially expressed genes in the T/NK cell cluster in both the 1 RFA and 3 RFA samples. **E)** Violin plot showing high Granzyme B mRNA levels in T/NK cluster following 1 and 3 RFA sessions**. F)** Violin plot showing significantly decreased Pdcd1 mRNA levels in the T/NK cell cluster (*p=0.027701) following 3 RFA sessions**. G)** Violin plot showing significantly increased Nt5e mRNA levels in the tumor clusters (*p=0.01859) and significantly decreased levels in the Csf1r+ macrophage cluster (*p=0.044041) following 3 RFA sessions**. H)** Violin plot showing high Cd274 mRNA levels in both the Il1b+ and Ccl3+ neutrophil clusters following 1 and 3 RFA sessions.

The tumor subclusters displayed heterogeneous phenotypes, characterized with distinct gene expression programs (**Fig. 2B, Supplemental Fig. S5C**). Cluster 0, which was decreased after three serial ablations, showed high expression of genes such as *Maff, Net1,* and *Sh3pdx2a*, which are characteristic of a stress-responsive invasive tumor cell state^36–38^. Cluster 3, which was increased after three serial ablations, had high gene expression of *Malat1* and *Ngf* which are characteristic of a mesenchymal/EMT-like tumor state which is an aggressive subtype driving perineural invasion^39,40^. Cluster 6, which was decreased after serial ablation, is a secretory/mesothelial-like tumor subtype characterized by high gene expression of *Msln* and *Mfge8*^41,42^. Cluster 8, which was increased after serial ablation, is a metabolically active tumor cell population, marked by gene expression of *Sgpp2* and *Degs2* which are key molecules in lipid metabolism and synthesis^43^. Lastly, cluster 9 was decreased following serial ablation and is a proliferative tumor cell cluster characterized by *Mik67* and cell cycle gene expression^44^. We assessed the expression of gene signatures associated with Classical, Exocrine, and Quasimesenchymal tumor phenotypes^45^. Across all tumor clusters, Quasimesenchymal, Exocrine and Classical signatures were identified; however, the Classical signature genes were most highly expressed after serial RFA treatments (**Fig. 2C**).

Given the experimental evidence of a more pronounced abscopal effect with serial RFA treatments, we next examined cytotoxic immune signatures within the T/NK cell cluster in RFA ablated tumors. We analyzed differentially expressed genes related to T cell identity, NK cell identity, and their activation/exhaustion states (**Fig. 2D**). In the single RFA-ablated tumors, there were very few or no cells expressing NK cell genes (*Nkg7, Klrb1c, Ncr1*) and there were both Cd4+ and Cd8b1+ T cells, however the Cd8+ T cells expressed low levels of Granzyme B (Gzmb) (**Fig. 2D and E**) and significantly higher levels of PD-1 (*Pdcd1*) (**Fig. 2D and F**) signaling a less active and more exhausted T cell profile compared to serially ablated tumors. Comparatively, in the serially ablated tumors, there were more abundant NK cells, as well as T cells expressing more *Gzmb*. Together, these results suggest there is enhanced cytotoxic immune activation after serial RFA treatments.

In addition to cytotoxic mediators, we assessed inflammatory cytokine expression across multiple cell types. Gene expression analysis revealed significantly elevated *Il18* mRNA expression following serial RFA treatments in the tumor cells, and expression in the macrophage, neutrophil, T/NK, and classical dendritic cell clusters (**Supplemental Fig. S5D**). This broad increase in *Il18* expression suggests serial RFA promotes a pro-inflammatory signaling environment and NK and T cell cytotoxic phenotypes across both immune and tumor compartments. To further characterize the TIM, we performed IHC staining for NK cells using NK1.1 (**Supplemental Fig. S5E**). Quantification revealed NK1.1^+^ cells were significantly increased in three RFA ablated and contralateral tumors compared to respective tumors from mice receiving one RFA treatment (**Supplemental Fig. S5F**). Furthermore, NK cell infiltration was significantly higher in the contralateral tumors from the three RFA-treated group compared with their RFA-treated counterparts.

Additionally, we assessed the innate immune infiltration of neutrophils, and found two distinct neutrophil clusters (**Supplemental Fig. S5G**). The first cluster is characterized by high expression of *Il1b* and the second cluster is characterized by high expression of *Ccl3* (**Supplemental Fig. S5H**). Both Il1b+ and Ccl3+ neutrophils have been shown to be pro-tumor associated populations of neutrophils^46,47^.

Previously, we have shown that RFA treatment increases intratumoral immune suppressive adenosine and that pharmacological inhibition of CD73, the ectoenzyme responsible for extracellular adenosine production, in combination with RFA treatment enhanced and sustained the RFA treatment effect^15^. Therefore, we analyzed changes in gene expression of *Nt5e*, the gene encoding for CD73, in response to serial RFA treatments. We found that while in the Csf1r+ macrophage cluster *Nt5e* expression was significantly decreased, in the tumor cluster the expression was significantly increased in response to serial RFA (**Fig. 2G**). We also have previously shown that RFA treated tumors increased the expression of PD-L1 compared to control tumors and the combination of RFA with anti-PD-L1 blockade inhibited tumor growth^14^. Therefore, we also analyzed gene expression of *Cd274* in response to serial ablations. We found widespread expression of PD-L1 in both the one RFA and three RFA ablated tumors. These data support the potential combination therapy targeting CD73 and the PD-1/PD-L1 axis in combination with serial RFA treatments.

### Serial RFA thermal ablation significantly enriches CSF1/CSF1R myeloid pathway signaling in ablated tumors

Proteomic profiling of tumor lysates further demonstrated serial ablation induces a robust inflammatory response. Tumors treated with three RFA sessions exhibited upregulation of multiple cytokines (**Fig. 3C**), including CSF1 (M-CSF), IFN-γ and TNF-α. IFN-γ and TNF-α are key cytokines that promote cytotoxic T cell activity through upregulation of ICAM-1 and enhanced antigen presentation to promote T cell inflammatory cell death and killing efficacy as well as promote the recruitment CD8^+^ T cells and NK cells^48–51^. However, many of the significantly increased cytokines are also key regulators of myeloid cell recruitment and differentiation during inflammatory tissue repair, including CSF1, LIF and GM-CSF, suggesting serial thermal injury also initiates a wound-healing-like response within the TME.

**Figure 3.**
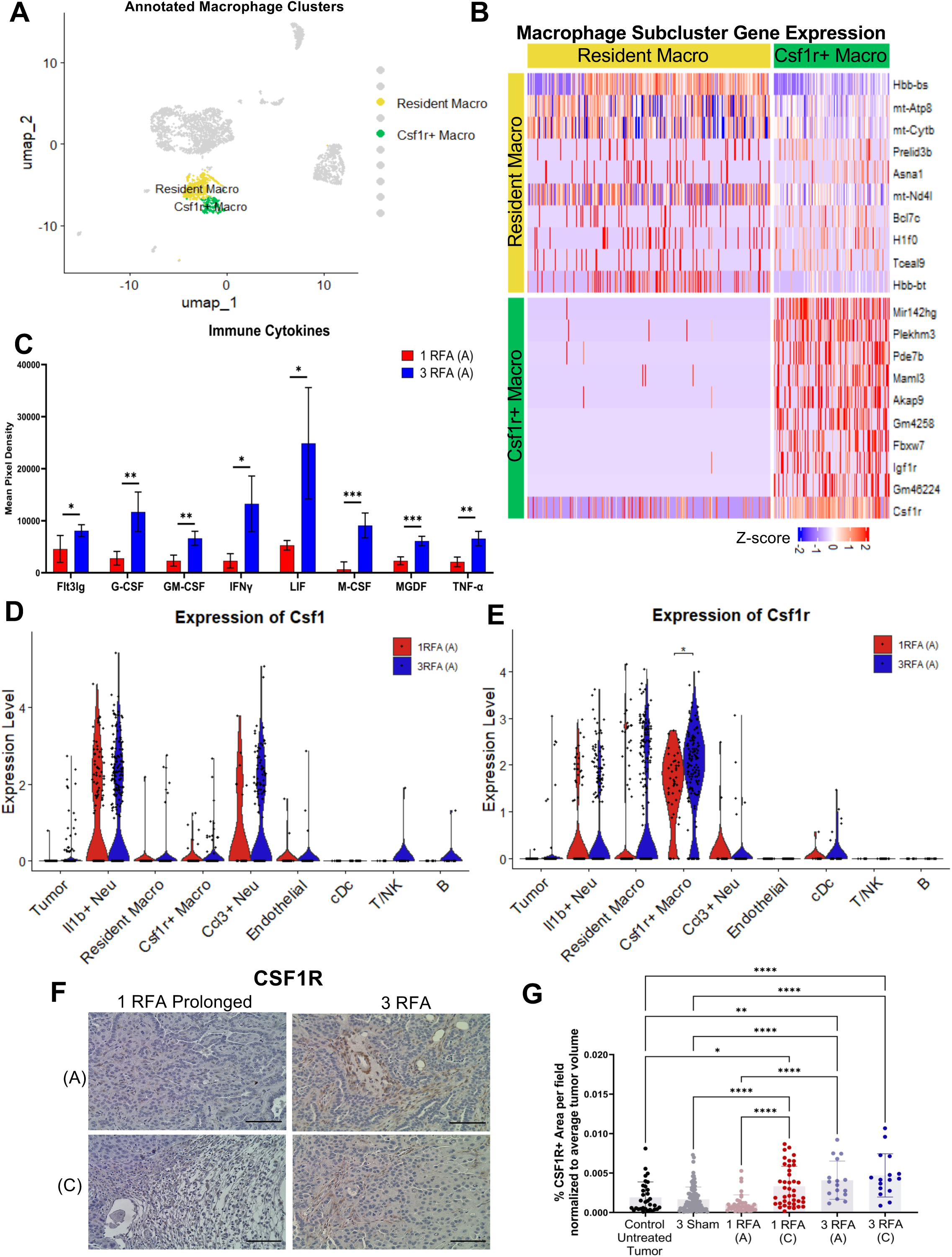
Serial RFA thermal ablation significantly enriches CSF1/CSF1R myeloid pathway signaling in ablated tumors. **A)** uMAP of 2 macrophage cell clusters from single-cell RNA-sequencing analysis. **B)** Heat map showing top differentially expressed genes in the 2 macrophage clusters. **C)** Proteome profiler shows Flt3lg (*p=0.0485), G-CSF (**p=0.0045), GM-CSF (**p=0.0026), IFNy (*p=0.0075), LIF (*p=0.0107), M-CSF (***p=0.0009), MDGF (***p=0.0007), and TNF-α (**p=0.0018) were all significantly increased immune cytokines in 3 RFA ablated tumors compared to 1 RFA ablated tumors. **D)** Violin plot of *Csf1* gene expression mRNA levels. **E)** Violin plot of *Csf1r* gene expression. *Csf1r* mRNA expression is significantly increased in the Csf1r+ macrophage cluster in three RFA-treated tumors (*p=0.031128). **F)** Representative 20x CSF1R immunohistochemistry images. Scale bars 50 µm. **G)** CSF1R staining is significantly increased in 3 RFA-treated and contralateral tumors compared to control, sham and 1 RFA-treated tumors (****p<0.0001). Csfr1 staining is significantly increased in 1 RFA contralateral tumors compared to control untreated (*p=0.0473) and 3 sham tumors (*p=0.0288). For **C, D, E,** and **G** an unpaired t-test was used for statistical analysis.

Consistent with this injury-driven response, in addition to increased CSF1 protein expression identified by the proteomic assay, scRNA-seq analysis demonstrated significant induction of *Csf1* expression across multiple immune populations in both the one and three RFA-treated tumors (**Fig. 3D**). Elevated *Csf1* transcripts were detected in both neutrophil clusters, as well as in the T/NK cell cluster and B cell cluster in serially treated tumors, indicating both innate and adaptive immune compartments contribute to CSF1 production in response to tumor injury. In parallel, expression of *Csf1r* was significantly increased within a macrophage subset characterized by low mitochondrial gene expression (**Fig. 3E**), consistent with expansion or activation of CSF1R⁺ macrophages. Activation of the CSF1/CSF1R signaling axis is a central component of wound-healing responses and promotes the recruitment, survival, and polarization of immune suppressive macrophages^52^. In tumor contexts, CSF1R⁺ myeloid cells are known to suppress cytotoxic lymphocyte function. Thus, while serial RFA promotes infiltration and activation of cytotoxic NK and T cells, it simultaneously induces a countervailing CSF1R-driven myeloid response that may restrain lymphocyte-mediated tumor killing.

Serial ablation also induced metabolic factor, stromal and extracellular matrix factor, growth factor, chemokine, and inflammation changes (**Supplemental Fig. S5A-F**). Proteome profiler analysis identified increased levels of proteins associated with extracellular matrix remodeling and inflammatory tissue repair (**Supplemental Fig. S5B**), including Tissue Factor (CD142), Dickkopf-related protein 1 (DKK-1), and Osteoprotegerin (OPG). Multiplex cytokine analysis further revealed increased levels of several interleukin family cytokines following three RFA sessions (**Supplemental Fig. S5G**). Additionally, multiplex cytokine analysis revealed serial ablation did not increase levels of IL-1b, IL-6, CXCL9, CXCl10, and increased levels of IL-12-40, all of which have been shown to be commonly secreted by M1-like tumor associated macrophages (**Supplemental Fig. S5**)^53,54,55^. Alternatively, CCL17 and CCL22, which are typically secreted by M2-like tumor associated macrophages^53,55^ was dramatically increased in three RFA-treated tumors (**Supplemental Fig. S5**). Collectively, these data demonstrate serial RFA reshapes the TME through an injury-induced inflammatory program characterized by increased cytotoxic lymphocyte infiltration together with a concurrent activation of CSF1/CSF1R-dependent myeloid pathways.

Additionally, immunohistochemical staining for CSF1R revealed significantly increased expression in three RFA-ablated tumors and contralateral tumors as well as single RFA-treated tumors compared to untreated tumors and tumors receiving three sham treatments (**Fig. 3F and G**).

### Serial thermal ablation in combination with anti-PD-L1 and Quemliclustat significantly reduces the volume of contralateral tumors

Preclinical studies have evaluated whether anti-PD-1/PD-L1 or targeting CD73 with Quemliclustat (Quemli) can enhance the tumor growth restraining capability of one RFA ablation^13–15^. We previously found the combination of Quemli with one RFA ablation significantly increased cytotoxic CD8^+^ T cell recruitment into ablated tumors^15^. We next investigated whether serial thermal ablation could be leveraged to enhance systemic antitumor immunity when combined with immunomodulatory therapies. Mice bearing bilateral tumors were treated with serial RFA of the primary tumor in combination with anti-PD-L1 and Quemli (**Fig. 4A**).

**Figure 4.**
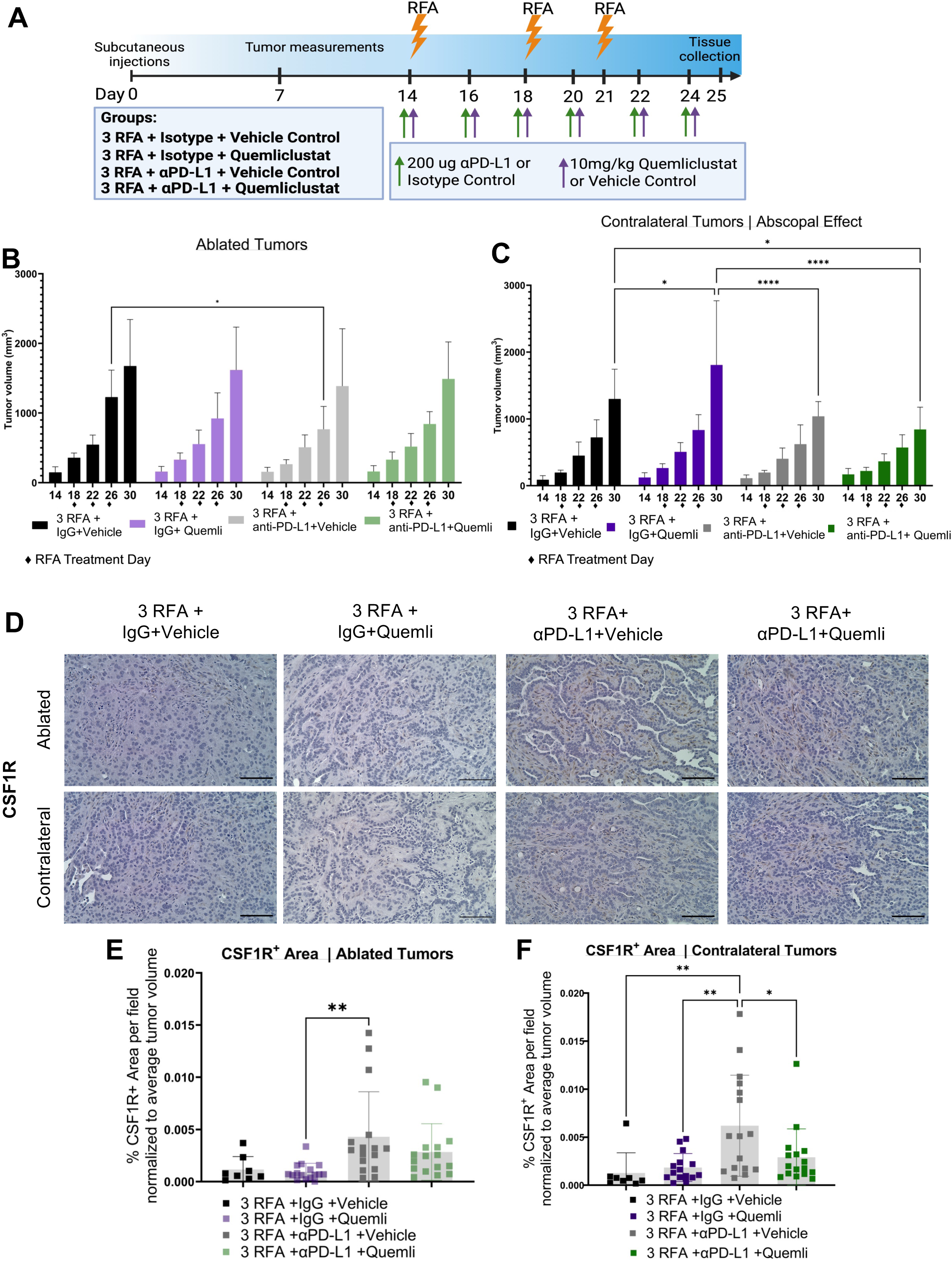
Repeated thermal ablation in combination with anti-PD-L1 and Quemliclustat significantly reduces the volume of contralateral tumors. **A**) Experimental design created using Biorender. **B)** Repeated thermal ablation using RFA in combination with anti-PD-L1 and Quemli does not significantly reduce the volume of treated tumors. **C)** However, repeated RFA in combination with anti-PD-L1 and Quemli significantly reduces the tumor volume of contralateral tumors compared to repeated RFA + IgG+Vehicle (p<0.05) and compared to repeated RFA + Quemli treated alone (p<0.0001). **D)** Representative 20x CSF1R immunohistochemistry images. Scale bars 100 μm. **E)** CSF1R+ cells are significantly increased in 3 RFA+anti-PD-L1+vehicle compared to 3 RFA+Quemli+IgG in the ablated tumors (**p=0.007). **F)** CSF1R+ cells are significantly increased in 3 RFA+anti-PD-L1+vehicle compared to 3 RFA+IgG+vehicle (**p=0.0095), 3 RFA+Quemli+IgG (**p=0.005), and 3 RFA+anti-PD-L1+Quemli (*p=0.0455) in the contralateral tumors. n=15 per group. A one-way Anova was used in Prism Graphpad for statistical comparisons.

Serial RFA combined with anti-PD-L1 and Quemli did not reduce the volume of the ablated tumors compared with control conditions (**Fig. 4B**); however, this therapeutic combination did produce a pronounced effect on distant contralateral tumor growth (**Fig. 4C**). Specifically, mice receiving serial RFA treatments together with anti-PD-L1 and Quemli exhibited a significant reduction in the growth of contralateral tumors relative to animals treated with serial RFA plus IgG and vehicle, serial RFA and anti-PD-L1, as well as those treated with serial RFA and Quemli alone (**Fig. 4C**), but did not significantly decrease the growth compared to contralateral tumors in the serial RFA-treated mice with anti-PD-L1.

We next examined the abundance of CSF1R^+^ cells within the TME of mice treated with serial RFA in combination with anti-PD-L1 and Quemli to determine if CSF1R^+^ cells increase in response to immune checkpoint blockade and serial RFA. Quantification of CSF1R^+^ staining revealed significantly increased CSF1R^+^ area per field in tumors treated with serial RFA in combination with anti-PD-L1, compared to tumors treated with three RFA and Quemli (**Fig. 4D-F**), indicating enhanced infiltration or expansion of CSF1R^+^ myeloid cells following these treatments. A more pronounced significant increase in CSF1R^+^ cell area was detected in the contralateral tumors where the three RFA-treated with Quemli and anti-PD-L1 had significantly more CSF1R+ cells per tumor volume than all other groups (**Fig. 4F**), suggesting systemic immune modulation induced by serial RFA treatment plus combination immunotherapy is associated with recruitment or accumulation of these myeloid populations at distant tumor sites.

### Serial thermal ablation in combination with CSF1R inhibition, anti-PD-L1 and Quemliclustat significantly reduces the volume of ablated and contralateral tumors

Given the observed activation of the CSF1/CSF1R signaling axis following serial RFA treatment, we next investigated whether targeting this myeloid compartment could further enhance therapeutic efficacy. So, we performed an additional experiment incorporating pharmacologic inhibition of CSF1R alongside anti-PD-L1 and Quemli during serial RFA treatment (**Fig. 5A)**. The addition of CSF1R inhibition in the treatment regimen resulted in a pronounced reduction in tumor growth at both the ablated and distant tumor sites compared to mice that received serial RFA with Quemli and anti-PD-L1 (**Fig. 5B-C**). Immune profiling of tumors from these mice revealed a significant increase in CD8^+^ and GZMB^+^ cells in ablated and contralateral tumors from mice that received three RFA treatments plus anti-PD-L1, Quemli and CSF1R inhibition compared to three RFA + anti-PD-L1 and Quemli (**Fig. 5D-I**). Together, these results demonstrate while serial thermal ablation combined with PD-L1 blockade and CD73 inhibition can promote systemic tumor control, the addition of CSF1R inhibition further improves therapeutic efficacy with a correlative increase in CD8^+^ and GZMB^+^ cells in both ablated and contralateral tumors.

**Figure 5.**
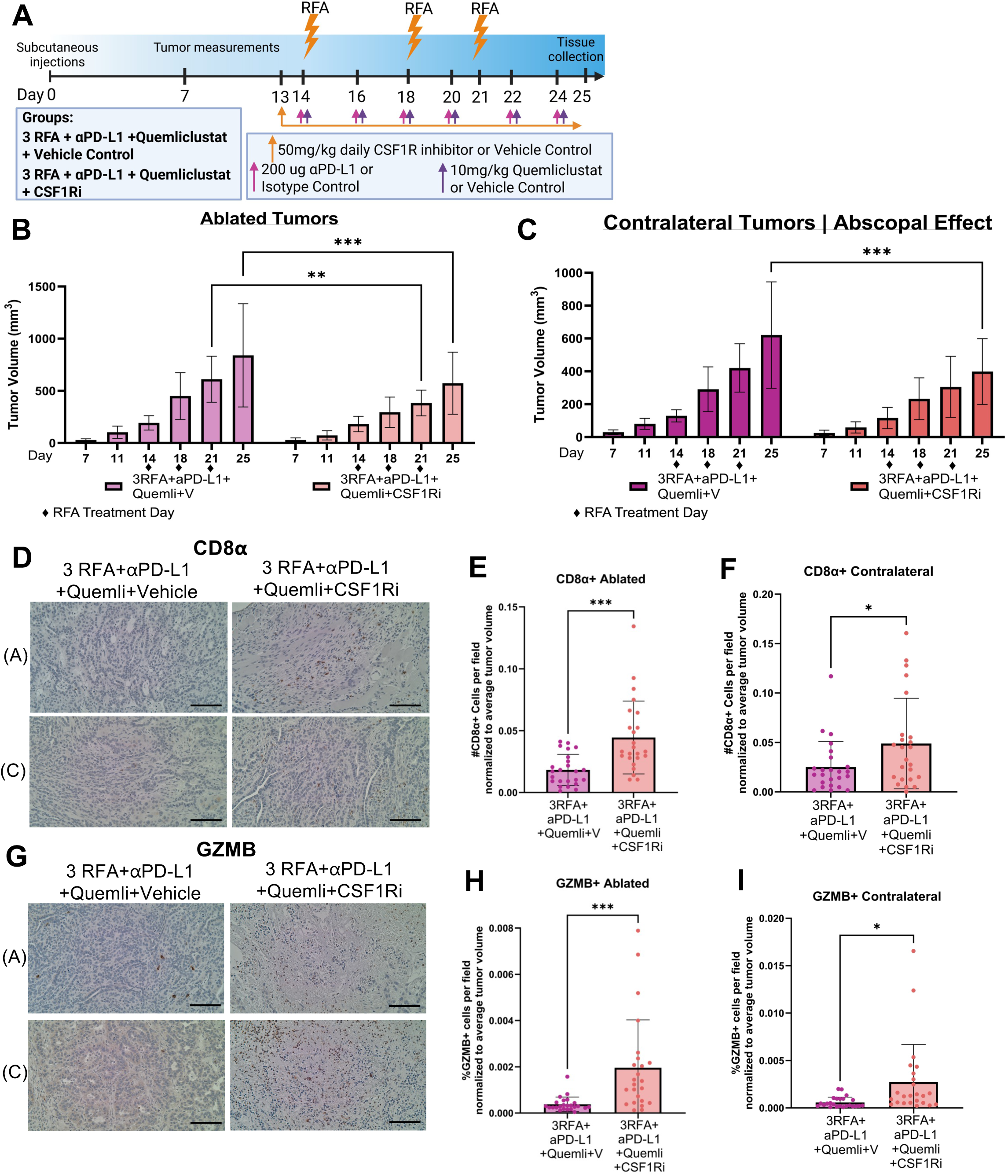
Serial thermal ablation in combination with CSF1R inhibition, anti-PD-L1 and Quemliclustat significantly reduces the volume of ablated and contralateral tumors. **A)** Experimental design created using biorender**. B)** Serial thermal ablation using RFA in combination with CSF1R inhibition, anti-PD-L1 and Quemli significantly reduces the volume of ablated tumors (**p<0.01; ***p<0.001) and **C)** contralateral tumors (**p<0.01; ***p<0.001). **D)** Representative 20x CD8α immunohistochemistry images. **E)** Combination treatment using 3 RFA+anti-PD-L1, Quemli and CSF1R inhibition significantly increases the infiltration of CD8α+ T cells in ablated (***p=0.0002) and **F)** contralateral tumors (*p=0.0318). **G)** Representative 20x Gzmb immunohistochemistry images. **H)** Combination treatment using 3 RFA+anti-PD-L1, Quemli and CSF1R inhibition significantly increases the infiltration of Gzmb+ T cells in ablated (***p=0.0006) and **I)** contralateral tumors (*p=0.0112). Scale bars 100 μm. n=15 per group. An unpaired t test was used for statistical comparisons.

### Serial thermal EUS-RFA enhances antitumor immune infiltration in long term survivors treated with EUS-RFA prior to surgical resection

To study the effects of serial thermal ablation on the tumor immune microenvironment in human PDAC patients, we stained samples from patients in the PANCARDINAL-1 clinical trial that received EUS-RFA and systemic chemotherapy prior to surgical resection. Patients who are long-term survivors (survival of greater than 4 years) had significantly increased infiltration of CD8α+ T cells compared to patients which did not receive EUS-RFA or short-term survivors (survival of less than 13 months) (**Fig. 6A-B**). Additionally, long term survivors had significantly decreased CD163+ immune suppressive macrophage infiltration compared to patients who did not receive EUS-RFA and short-term survivors (**Fig. 6C-D**). Together, these data suggest serial EUS-RFA thermal ablations reshaped the tumor immune microenvironment to an antitumor immune landscape, correlating with prolonged survival.

**Figure 6.**
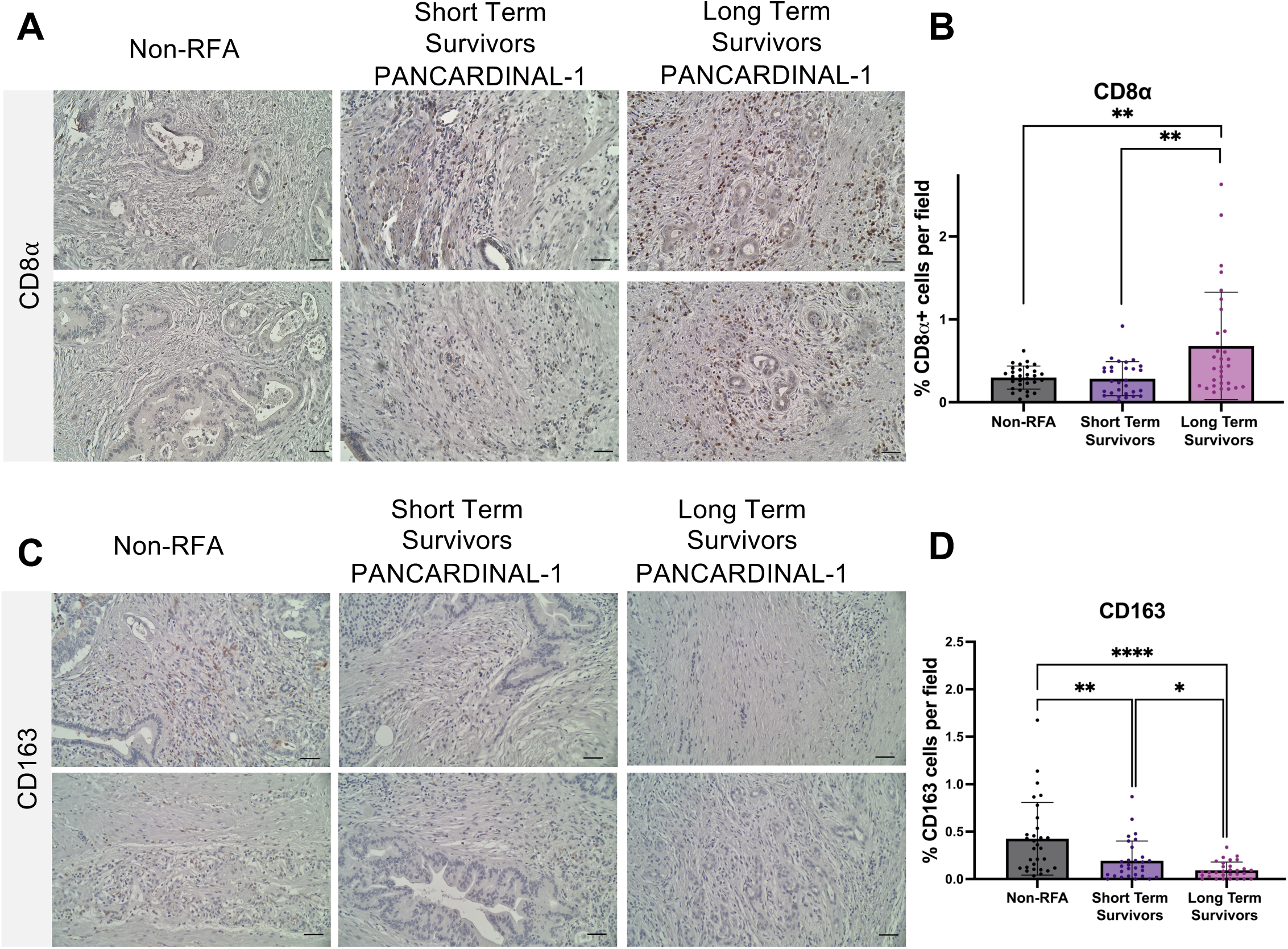
Long-term survivors after EUS-RFA show significantly increased CD8α^+^ cells. **A)** Representative 20x images of CD8a staining in non-RFA PDAC patients, Long-term PDAC survivors (greater than 4 years) treated with at least 3 EUS-RFA procedures prior to surgery, and Short-term survivors (less than 13 months survival) expression treated with 1-3 EUS-RFA procedures prior to surgery. **B)** CD8α^+^ cells are significantly increased in Long-term survivors compared to non-RFA patients (**p=0.0026) and Short-term survivors (**p=0.0027). **C)** Representative 20x images of CD163 staining in non-RFA PDAC patients, Long-term PDAC survivors (greater than 4 years) treated with at least 3 EUS-RFA procedures prior to surgery, and Short-term survivors (less than 13 months survival) expression treated with 1-3 EUS-RFA procedures prior to surgery. **D)** CD163+ cells are significantly decreased in Short-term survivors compared to non-RFA patients (*p=0.0050). CD163+ cells are significantly decreased in Long-term survivors compared to non-RFA patients (****p<0.0001) and Short-term survivors (*p=0.0179). n=3 clinical samples per group. Scale bars 50uM.

## DISCUSSION

Targeted thermal ablation is increasingly recognized not only as a local cytoreductive therapy but also as a stimulus for systemic immune activation within the tumor microenvironment. In this study, we demonstrate serial RFA treatment produces substantially greater antitumor effects than one ablation, enhancing necrosis, cytotoxic immune responses, and growth inhibition in both treated and contralateral tumors. Serial ablation reshapes the immune landscape, including expansion of neutrophils and dynamic alterations in macrophage populations, highlighting the role of innate immunity in mediating both local and systemic responses. Single-cell transcriptomic analysis further identifies a CSF1-driven myeloid program that may limit the durability of these responses, revealing a targetable mechanism of adaptive resistance.

Emerging clinical data reinforce the translational relevance of immunomodulatory combinations in PDAC. The phase 1b ARC-8 trial evaluated Quemliclustat combined with anti-PD-1 therapy and chemotherapy in patients with advanced PDAC. Post-treatment analyses linked maximal suppression of adenosine-associated transcripts, including the *NR4A* gene family, with enhanced T-cell activation and survival benefit^56^. Similarly, a phase 2 study combining focal radiation with dual checkpoint blockade in microsatellite-stable colorectal and pancreatic cancer demonstrated local irradiation can sensitize tumors otherwise resistant to immunotherapy, producing disease control in a subset of patients^57^. Retrospective and prospective analyses of stereotactic body radiotherapy, irreversible electroporation, RFA and other targeted ablative modalities combined with systemic therapy across solid tumors have further suggested timing, sequencing, and frequency of local interventions critically influence immune priming and clinical outcomes^58,59^. These clinical observations support the concept that immune responses to tumor-directed injury are dynamic and potentially programmable.

Our preclinical data provide mechanistic insight into these clinical findings. Serial RFA amplifies systemic cytotoxic immunity but also induces compensatory expansion of Ccl3^+^ neutrophils, recently defined as pro-tumor^47^ and CSF1R^+^ regulatory myeloid populations. These results suggest the interval between ablation events and the timing of combination immunotherapy will determine the balance between immune activation and adaptive suppression. Optimizing the temporal relationship between tumor injury and systemic therapy could enhance the durability of antitumor responses and improve clinical outcomes.

In addition to innate immune remodeling, serial RFA induces a broad inflammatory transcriptional program characterized by increased *Il18* expression across multiple cell types. IL-18 is a cytokine that promotes natural killer cell activation, enhances T-cell–mediated immunity, and can augment immunotherapy responses, indicating serial ablation generates a cytokine environment favorable for sustained cytotoxic activity. Pathway analysis revealed activation of T-cell receptor, NK cell, and Th1-associated signaling within infiltrating lymphocytes following serial RFA, accompanied by reduced PD-1/PD-L1 pathway activity. These findings highlight how iterative tumor injury can prime the immune system while also eliciting regulatory mechanisms that may constrain the response.

Finally, evaluation of patient samples treated with serial EUS-RFA in combination with systemic chemotherapy revealed significant increases in CD8+ T cell infiltration and significantly fewer CD163+ immune suppressive macrophages in long-term survivors compared with short-term survivors and non-RFA treated patients. These observations indicate that the combination of serial EUS-RFA and systemic chemotherapy remodels the tumor immune microenvironment to an anti-tumor phenotype. Although the decrease in immune suppressive macrophages is contrary to our findings in the animal model, this finding is consistent with studies showing that chemotherapy regimens are capable of reducing immune suppressive macrophages in the PDAC microenvironment^60,61^.

Together, these findings define targetable resistance mechanisms and provide a rationale for combinatorial strategies integrating serial tumor ablation with systemic immunotherapy. Rational scheduling of ablation events, consideration of immunosuppressive feedback, and integration of checkpoint or adenosine-targeted therapies represent actionable strategies to convert immunologically cold tumors into responsive disease. Future studies should prospectively evaluate how timing, dose, and sequence of ablation influence immune priming, as well as the functional contributions of neutrophils and CSF1R-positive myeloid cells, to fully translate these insights into effective clinical interventions in PDAC.

## ACKNOWLEDGEMENTS

WL was a CPRIT Predoctoral Fellow in the Biomedical Informatics, Genomics and Translational Cancer Research Training Program (BIG-TCR) funded by Cancer Prevention & Research Institute of Texas (CPRIT RP210045). We thank the technical support from UTHealth Cancer Genomics Core funded by the CPRIT grant (RP240610). Research reported in this publication was supported by the UNMC Pancreatic Cancer Center of Excellence and National Cancer Institute of the National Institutes of Health under award number P30 CA036727 (UNMC CCSG Cancer Center Grant). The content is solely the responsibility of the authors and does not necessarily represent the official views of the National Institutes of Health.

## Data Transparency Statement

All the raw and processed data supporting the findings of this study are available through GEO, accession number GSE325036. The R scripts supporting the findings of this paper are available upon request.

## Author contributions

LNS: Conceptualization, Data curation, Formal analysis, Writing-original draft, MVD: Data curation, Formal analysis, WL: Data curation, Formal analysis, AMW: Data curation, SD: Data curation, KT: Data curation, UKS: Writing-review and editing, NRM: Data curation, CVK: Data curation, BO: Data curation, JR, Writing-review and editing, PC: Investigation, KAK: Formal analysis, JLC: Formal analysis, ZZ: Funding acquisition, Formal analysis, SRH: Conceptualization, Formal analysis, Writing-review and editing, CJW: Formal analysis, Writing-review and editing, NCT: Conceptualization, Funding acquisition, JMB-L: Conceptualization, Formal analysis, Funding acquisition, Supervision, Writing-original draft

## REFERENCES

1. Lemdani K, Mignet N, Boudy V, et al. Local immunomodulation combined to radiofrequency ablation results in a complete cure of local and distant colorectal carcinoma. Oncoimmunology. 2019;8(3):1550342. doi:10.1080/2162402X.2018.1550342

2. Scopelliti F, Pea A, Conigliaro R, et al. Technique, safety, and feasibility of EUS-guided radiofrequency ablation in unresectable pancreatic cancer. Surg Endosc. Sep 2018;32(9):4022–4028. doi:10.1007/s00464-018-6217-x

3. Goyal D CP, Wray CJ, et al. 1111 Feasibility, Safety, and Efficacy of Endoscopic Ultrasound (EUS) Guided Radiofrequency Ablation (RFA) of the Pancreatic Lesions: Single Center Us Experience. Gastrointestinal Endoscopy 2017.

4. Song TJ, Seo DW, Lakhtakia S, et al. Initial experience of EUS-guided radiofrequency ablation of unresectable pancreatic cancer. Gastrointest Endosc. Feb 2016;83(2):440–3. doi:10.1016/j.gie.2015.08.048

5. Pai M, Habib N, Senturk H, et al. Endoscopic ultrasound guided radiofrequency ablation, for pancreatic cystic neoplasms and neuroendocrine tumors. World J Gastrointest Surg. Apr 2015;7(4):52–9. doi:10.4240/wjgs.v7.i4.52

6. Lorber G, Glamore M, Doshi M, Jorda M, Morillo-Burgos G, Leveillee RJ. Long-term oncologic outcomes following radiofrequency ablation with real-time temperature monitoring for T1a renal cell cancer. Urol Oncol. Oct 2014;32(7):1017–23. doi:10.1016/j.urolonc.2014.03.005

7. Girelli R, Frigerio I, Giardino A, et al. Results of 100 pancreatic radiofrequency ablations in the context of a multimodal strategy for stage III ductal adenocarcinoma. Langenbecks Arch Surg. Jan 2013;398(1):63–9. doi:10.1007/s00423-012-1011-z

8. Yokoyama T, Egami K, Miyamoto M, et al. Percutaneous and laparoscopic approaches of radiofrequency ablation treatment for liver cancer. J Hepatobiliary Pancreat Surg. 2003;10(6):425–7. doi:10.1007/s00534-002-0830-7

9. Curley SA, Izzo F, Ellis LM, Nicolas Vauthey J, Vallone P. Radiofrequency ablation of hepatocellular cancer in 110 patients with cirrhosis. Ann Surg. Sep 2000;232(3):381–91. doi:10.1097/00000658-200009000-00010

10. Crinò SF, D’Onofrio M, Bernardoni L, et al. EUS-guided Radiofrequency Ablation (EUS-RFA) of Solid Pancreatic Neoplasm Using an 18-gauge Needle Electrode: Feasibility, Safety, and Technical Success. J Gastrointestin Liver Dis. Mar 2018;27(1):67–72. doi:10.15403/jgld.2014.1121.271.eus

11. Lakhtakia S, Ramchandani M, Galasso D, et al. EUS-guided radiofrequency ablation for management of pancreatic insulinoma by using a novel needle electrode (with videos). Gastrointest Endosc. Jan 2016;83(1):234–9. doi:10.1016/j.gie.2015.08.085

12. Moond V, Maniyar B, Harne PS, Bailey-Lundberg JM, Thosani NC. Harnessing endoscopic ultrasound-guided radiofrequency ablation to reshape the pancreatic ductal adenocarcinoma microenvironment and elicit systemic immunomodulation. Explor Target Antitumor Ther. 2024;5(5):1056–1073. doi:10.37349/etat.2024.00263

13. Musiu C, Adamo A, Caligola S, et al. Local ablation disrupts immune evasion in pancreatic cancer. Cancer Lett. Jan 28 2025;609:217327. doi:10.1016/j.canlet.2024.217327

14. Faraoni EY, O’Brien BJ, Strickland LN, et al. Radiofrequency Ablation Remodels the Tumor Microenvironment and Promotes Neutrophil-Mediated Abscopal Immunomodulation in Pancreatic Cancer. Cancer Immunol Res. Jan 03 2023;11(1):4–12. doi:10.1158/2326-6066.CIR-22-0379

15. Faraoni EY, Strickland LN, O’Brien BJ, et al. Radiofrequency ablation in combination with CD73 inhibitor AB680 reduces tumor growth and enhances anti-tumor immunity in a syngeneic model of pancreatic ductal adenocarcinoma. Front Oncol. 2022;12:995027. doi:10.3389/fonc.2022.995027

16. Zerbini A, Pilli M, Penna A, et al. Radiofrequency thermal ablation of hepatocellular carcinoma liver nodules can activate and enhance tumor-specific T-cell responses. Cancer Res. Jan 15 2006;66(2):1139–46. doi:10.1158/0008-5472.CAN-05-2244

17. Dromi SA, Walsh MP, Herby S, et al. Radiofrequency ablation induces antigen-presenting cell infiltration and amplification of weak tumor-induced immunity. Radiology. Apr 2009;251(1):58–66. doi:10.1148/radiol.2511072175

18. Waitz R, Solomon SB, Petre EN, et al. Potent induction of tumor immunity by combining tumor cryoablation with anti-CTLA-4 therapy. Cancer Res. Jan 15 2012;72(2):430–9. doi:10.1158/0008-5472.CAN-11-1782

19. Shi L, Chen L, Wu C, et al. PD-1 Blockade Boosts Radiofrequency Ablation-Elicited Adaptive Immune Responses against Tumor. Clin Cancer Res. Mar 01 2016;22(5):1173–1184. doi:10.1158/1078-0432.CCR-15-1352

20. Liu X, Baer JM, Stone ML, et al. Stromal reprogramming overcomes resistance to RAS-MAPK inhibition to improve pancreas cancer responses to cytotoxic and immune therapy. Sci Transl Med. Oct 23 2024;16(770):eado2402. doi:10.1126/scitranslmed.ado2402

21. Lander VE, Belle JI, Kingston NL, et al. Stromal Reprogramming by FAK Inhibition Overcomes Radiation Resistance to Allow for Immune Priming and Response to Checkpoint Blockade. Cancer Discov. Dec 02 2022;12(12):2774–2799. doi:10.1158/2159-8290.CD-22-0192

22. Wray CJ, O’Brien B, Cen P, et al. EUS-guided radiofrequency ablation for pancreatic adenocarcinoma. Gastrointest Endosc. Oct 2024;100(4):759–766. doi:10.1016/j.gie.2024.04.2926

23. Thosani N, Cen P, Rowe J, et al. Endoscopic ultrasound-guided radiofrequency ablation (EUS-RFA) for advanced pancreatic and periampullary adenocarcinoma. Sci Rep. Oct 03 2022;12(1):16516. doi:10.1038/s41598-022-20316-2

24. Bianchi A, De Castro Silva I, Deshpande NU, et al. Cell-Autonomous Cxcl1 Sustains Tolerogenic Circuitries and Stromal Inflammation via Neutrophil-Derived TNF in Pancreatic Cancer. Cancer Discov. Jun 02 2023;13(6):1428–1453. doi:10.1158/2159-8290.CD-22-1046

25. Garcia Garcia CJ, Huang Y, Fuentes NR, et al. Stromal HIF2 Regulates Immune Suppression in the Pancreatic Cancer Microenvironment. Gastroenterology. Jun 2022;162(7):2018–2031. doi:10.1053/j.gastro.2022.02.024

26. Yang D, Liu J, Qian H, Zhuang Q. Cancer-associated fibroblasts: from basic science to anticancer therapy. Exp Mol Med. Jul 2023;55(7):1322–1332. doi:10.1038/s12276-023-01013-0

27. Zhu Y, Knolhoff BL, Meyer MA, et al. CSF1/CSF1R blockade reprograms tumor-infiltrating macrophages and improves response to T-cell checkpoint immunotherapy in pancreatic cancer models. Cancer Res. Sep 15 2014;74(18):5057–69. doi:10.1158/0008-5472.CAN-13-3723

28. Stromnes IM, Burrack AL, Hulbert A, et al. Differential Effects of Depleting versus Programming Tumor-Associated Macrophages on Engineered T Cells in Pancreatic Ductal Adenocarcinoma. Cancer Immunol Res. Jun 2019;7(6):977–989. doi:10.1158/2326-6066.CIR-18-0448

29. Candido JB, Morton JP, Bailey P, et al. CSF1R. Cell Rep. May 01 2018;23(5):1448–1460. doi:10.1016/j.celrep.2018.03.131

30. Hingorani SR. Epithelial and stromal co-evolution and complicity in pancreatic cancer. Nat Rev Cancer. Feb 2023;23(2):57–77. doi:10.1038/s41568-022-00530-w

31. Hao Y, Stuart T, Kowalski MH, et al. Dictionary learning for integrative, multimodal and scalable single-cell analysis. Nat Biotechnol. Feb 2024;42(2):293–304. doi:10.1038/s41587-023-01767-y

32. Germain PL, Lun A, Garcia Meixide C, Macnair W, Robinson MD. Doublet identification in single-cell sequencing data using. F1000Res. 2021;10:979. doi:10.12688/f1000research.73600.2

33. Korsunsky I, Millard N, Fan J, et al. Fast, sensitive and accurate integration of single-cell data with Harmony. Nat Methods. Dec 2019;16(12):1289–1296. doi:10.1038/s41592-019-0619-0

34. Gu Z, Eils R, Schlesner M. Complex heatmaps reveal patterns and correlations in multidimensional genomic data. Bioinformatics. Sep 15 2016;32(18):2847–9. doi:10.1093/bioinformatics/btw313

35. Wickham H. ggplot2. Use R! Springer Nature; 2016.

36. Todoric J, Antonucci L, Di Caro G, et al. Stress-Activated NRF2-MDM2 Cascade Controls Neoplastic Progression in Pancreas. Cancer Cell. Dec 11 2017;32(6):824–839.e8. doi:10.1016/j.ccell.2017.10.011

37. Carr HS, Zuo Y, Oh W, Frost JA. Regulation of focal adhesion kinase activation, breast cancer cell motility, and amoeboid invasion by the RhoA guanine nucleotide exchange factor Net1. Mol Cell Biol. Jul 2013;33(14):2773–86. doi:10.1128/MCB.00175-13

38. Xu D, He Y, Zhao P, Liao C, Tan J. Pan-cancer multi-omics dissection of podosome-related genes reveals their dual roles in tumor invasion and immune modulation. Discov Oncol. Feb 12 2026;17(1)doi:10.1007/s12672-026-04558-4

39. Cheng Y, Imanirad P, Jutooru I, et al. Role of metastasis-associated lung adenocarcinoma transcript-1 (MALAT-1) in pancreatic cancer. PLoS One. 2018;13(2):e0192264. doi:10.1371/journal.pone.0192264

40. Shen S, Wang Q, Wang X, et al. Nodal Enhances Perineural Invasion in Pancreatic Cancer by Promoting Tumor-Nerve Convergence. J Healthc Eng. 2022;2022:9658890. doi:10.1155/2022/9658890

41. Montemagno C, Cassim S, Pouyssegur J, Broisat A, Pagès G. From Malignant Progression to Therapeutic Targeting: Current Insights of Mesothelin in Pancreatic Ductal Adenocarcinoma. Int J Mol Sci. Jun 06 2020;21(11)doi:10.3390/ijms21114067

42. Gu Y, Zhang Z, Camps MGM, Ossendorp F, Wijdeven RH, Ten Dijke P. Genome-wide CRISPR screens define determinants of epithelial-mesenchymal transition mediated immune evasion by pancreatic cancer cells. Sci Adv. Jul 14 2023;9(28):eadf9915. doi:10.1126/sciadv.adf9915

43. Zhao L, Zhao H, Yan H. Gene expression profiling of 1200 pancreatic ductal adenocarcinoma reveals novel subtypes. BMC Cancer. May 29 2018;18(1):603. doi:10.1186/s12885-018-4546-8

44. Karamitopoulou E, Zlobec I, Tornillo L, et al. Differential cell cycle and proliferation marker expression in ductal pancreatic adenocarcinoma and pancreatic intraepithelial neoplasia (PanIN). Pathology. Apr 2010;42(3):229–34. doi:10.3109/00313021003631379

45. Collisson EA, Sadanandam A, Olson P, et al. Subtypes of pancreatic ductal adenocarcinoma and their differing responses to therapy. Nat Med. Apr 2011;17(4):500–3. doi:10.1038/nm.2344

46. Wang L, Liu Y, Dai Y, et al. Single-cell RNA-seq analysis reveals BHLHE40-driven pro-tumour neutrophils with hyperactivated glycolysis in pancreatic tumour microenvironment. Gut. May 2023;72(5):958–971. doi:10.1136/gutjnl-2021-326070

47. Bolli E, Wirapati P, Hicham M, et al. CCL3 is produced by aged neutrophils across cancers and promotes tumor growth. Cancer Cell. Mar 09 2026;44(3):624–640.e12. doi:10.1016/j.ccell.2026.01.006

48. Bhat P, Leggatt G, Waterhouse N, Frazer IH. Interferon-γ derived from cytotoxic lymphocytes directly enhances their motility and cytotoxicity. Cell Death Dis. Jun 01 2017;8(6):e2836. doi:10.1038/cddis.2017.67

49. Wang R, Jaw JJ, Stutzman NC, Zou Z, Sun PD. Natural killer cell-produced IFN-γ and TNF-α induce target cell cytolysis through up-regulation of ICAM-1. J Leukoc Biol. Feb 2012;91(2):299–309. doi:10.1189/jlb.0611308

50. Karki R, Sharma BR, Tuladhar S, et al. Synergism of TNF-α and IFN-γ Triggers Inflammatory Cell Death, Tissue Damage, and Mortality in SARS-CoV-2 Infection and Cytokine Shock Syndromes. Cell. Jan 07 2021;184(1):149–168.e17. doi:10.1016/j.cell.2020.11.025

51. Barberini F, Pietroni R, Ielpo S, et al. Combined IFN-γ and TNF-α treatment enhances the susceptibility of breast cancer cells and spheroids to Natural Killer cell-mediated killing. Cell Death Dis. Oct 16 2025;16(1):729. doi:10.1038/s41419-025-08021-0

52. Sun L, Clavijo PE, Robbins Y, et al. Inhibiting myeloid-derived suppressor cell trafficking enhances T cell immunotherapy. JCI Insight. Apr 04 2019;4(7)doi:10.1172/jci.insight.126853

53. Ostuni R, Kratochvill F, Murray PJ, Natoli G. Macrophages and cancer: from mechanisms to therapeutic implications. Trends Immunol. Apr 2015;36(4):229–39. doi:10.1016/j.it.2015.02.004

54. Cannarile MA, Weisser M, Jacob W, Jegg AM, Ries CH, Rüttinger D. Colony-stimulating factor 1 receptor (CSF1R) inhibitors in cancer therapy. J Immunother Cancer. Jul 18 2017;5(1):53. doi:10.1186/s40425-017-0257-y

55. Noy R, Pollard JW. Tumor-associated macrophages: from mechanisms to therapy. Immunity. Jul 17 2014;41(1):49–61. doi:10.1016/j.immuni.2014.06.010

56. Wainberg ZA, Manji GA, Bahary N, et al. Quemliclustat and chemotherapy with or without zimberelimab in metastatic pancreatic adenocarcinoma: a randomized phase 1 trial. Nat Med. Mar 30 2026;doi:10.1038/s41591-026-04283-z

57. Parikh AR, Szabolcs A, Allen JN, et al. Radiation therapy enhances immunotherapy response in microsatellite stable colorectal and pancreatic adenocarcinoma in a phase II trial. Nat Cancer. Nov 2021;2(11):1124–1135. doi:10.1038/s43018-021-00269-7

58. Woeste MR, Shrestha R, Geller AE, et al. Irreversible electroporation augments β-glucan induced trained innate immunity for the treatment of pancreatic ductal adenocarcinoma. J Immunother Cancer. Apr 2023;11(4)doi:10.1136/jitc-2022-006221

59. Granata V, Grassi R, Fusco R, et al. Local ablation of pancreatic tumors: State of the art and future perspectives. World J Gastroenterol. Jun 21 2021;27(23):3413–3428. doi:10.3748/wjg.v27.i23.3413

60. Pratt HG, Steinberger KJ, Mihalik NE, et al. Macrophage and Neutrophil Interactions in the Pancreatic Tumor Microenvironment Drive the Pathogenesis of Pancreatic Cancer. Cancers (Basel). Dec 31 2021;14(1)doi:10.3390/cancers14010194

61. Long Y, Xie L, Xie F, et al. Targeting tumor-associated macrophages in pancreatic cancer. ILIVER. Jun 2026;5(2):100243. doi:10.1016/j.iliver.2026.100243

