## Supplemental Figures for "Combination Immunotherapy Enhances Serial Radiofrequency Ablation Induced Systemic Antitumor Immunity in Pancreatic Cancer"

**Figure S1**

**A** Sample 1 (1RFA) Doublet Classification

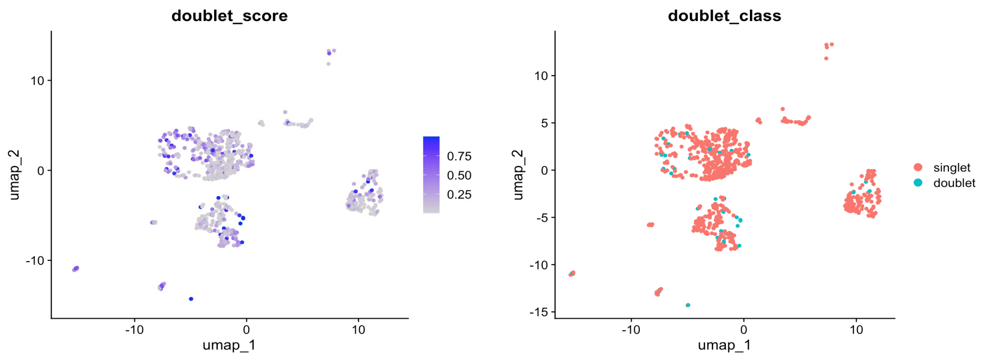


**B** Sample 2 (3RFA) Doublet Classification

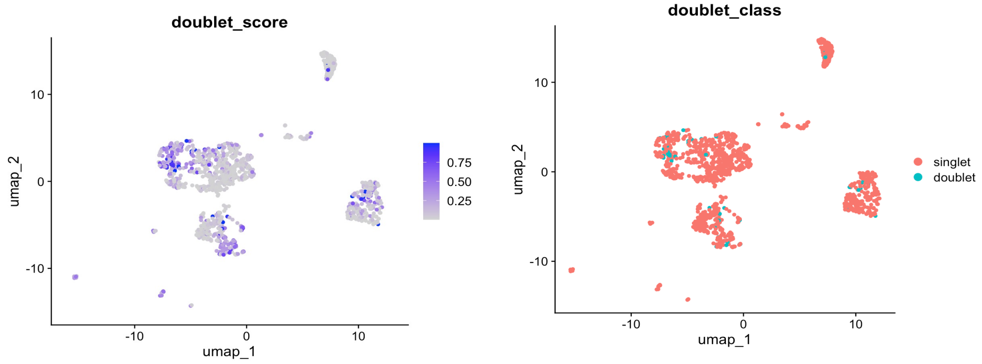


**C** Sample 3 (3RFA) Doublet Classification

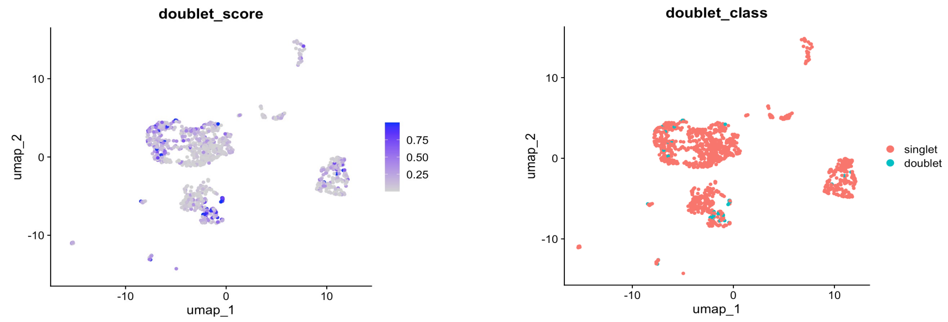


**D**

| **Sample** | **1** | **2** | **3** |
|---|---|---|---|
| Singlets | 728 | 1306 | 1186 |
| Doublets | 41 | 54 | 45 |
| % Doublets | 5.6 | 4.1 | 3.8 |

**E** Clustering without Doublets

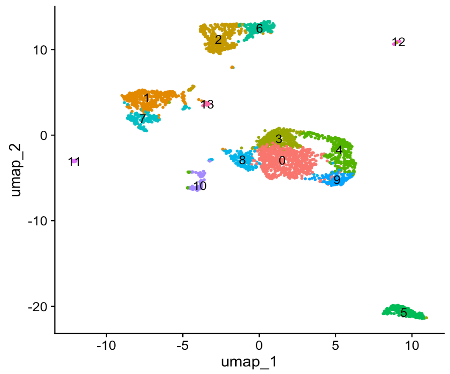


**F** Clustering Identification without Doublets

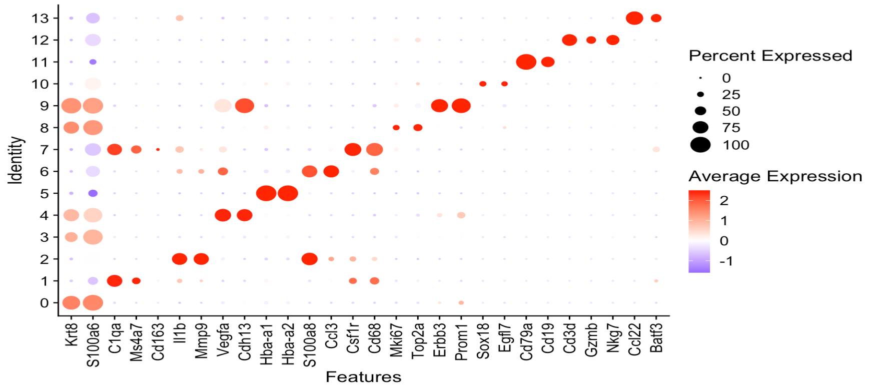


**Supplemental Figure S1. scRNA-seq doublet classification. A)** Sample 1 (1RFA) doublet classification. **B)** Sample 2 (3RFA) doublet classification. **C)** Sample 3 (3RFA) doublet classification. **D)** Table showing the %doublets per sample. **E)** uMAP of cell clustering without doublets. **F)** Clustering identification without doublets.

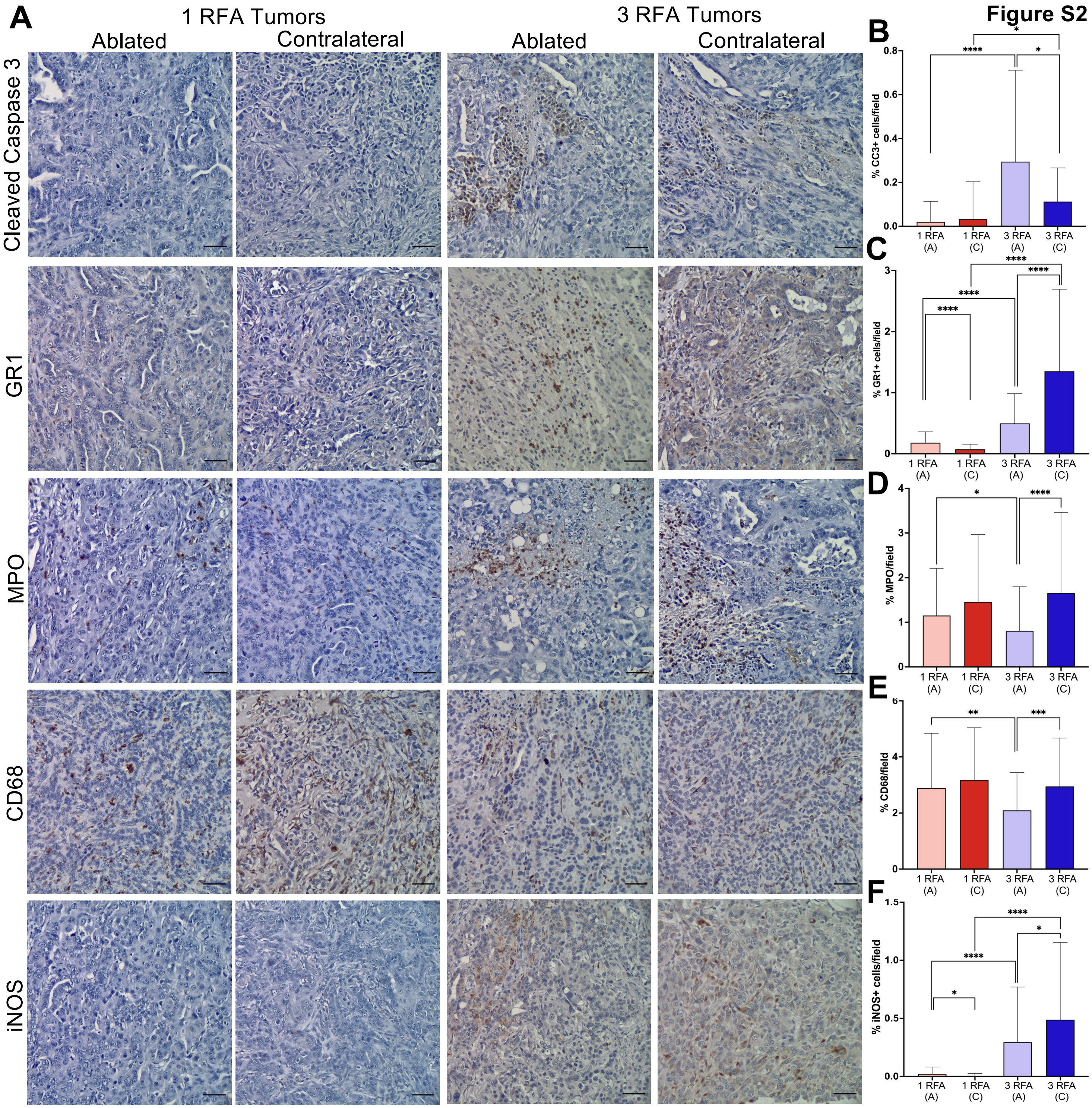


**Supplemental Figure S2. Serial RFA increases apoptosis and immune cell infiltration**. **A)** Representative 20x IHC images of CC3, GR1, MPO, CD68, and iNOS. Scale bars 50 μm. **B)** Serial thermal ablation significantly increases the abundance of cleaved caspase 3+ cells in both RFA-treated tumors (****p<0.0001) and contralateral tumors (*p=0.0436) compared to the respective 1 RFA tumors. CC3+ cells are also significantly increased in 3 RFA treated tumors compared to 3 RFA contralateral tumors (*p=0.0234). **C)** GR1 is significantly increased in 3 RFA RFA-treated tumors compared to 1 RFA treated tumors (****p<0.0001). GR1 is significantly increased in 3 RFA contralateral tumors compared to 1 RFA contralateral tumors (****p<0.0001). GR1 is also increased in 1 RFA treated tumors compared to 1 RFA contralateral tumors (****p<0.0001). GR1 is also increased in 3 RFA contralateral tumors compared to 3 RFA RFA-treated tumors (****p<0.0001). n=87 fields analyzed for 1 RFA tumors and n=109 fields analyzed for 3 RFA tumors. **D)** Serial RFA significantly increases the number of MPO+ cells per field in contralateral compared to RFA treated tumors (****p<0.0001). Serial RFA significantly reduces MPO levels in the ablated tumors compared to 1 RFA ablated tumors (*p=0.0253). **E)** Serial RFA significantly reduces the %CD68+ cells per field in RFA treated tumors compared to 1 RFA treated tumors (**p=0.0028) and 3 RFA contralateral tumors (***p=0.0003). **F)** iNOS is significantly increased in both the 3 RFA RFA-treated (****p<0.0001) and contralateral (****p<0.0001) tumors compared to the respective 1 RFA tumors. iNOS is significantly increased in 1 RFA treated tumors compared to 1 RFA contralateral tumors (*p=0.0432). iNOS is also increased in 3 RFA contralateral tumors compared to 3 RFA RFA-treated tumors (*p=0.0334). n=90 fields analyzed for 1 RFA tumors and n=82 fields analyzed for 3 RFA tumors. A student's t-test in Graphpad Prism was used for statistical comparisons.

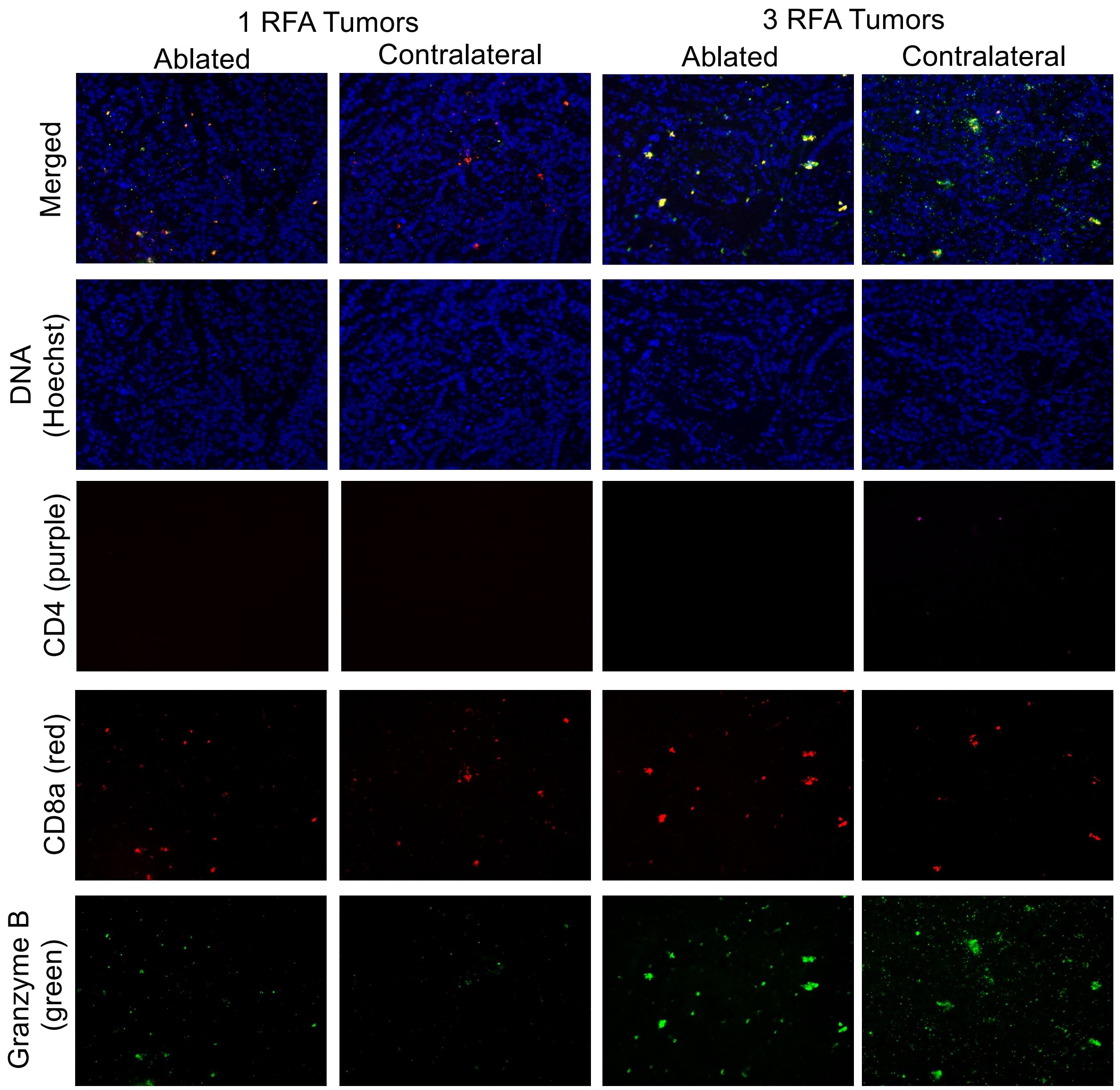


**Supplemental Figure S3. Repeated RFA increases infiltration of CD8α+ GZMB+ cells**. Individual panel IF from CD4, CD8 and granzyme B IF analysis of 1 and 3 RFA treated and contralateral tumors.

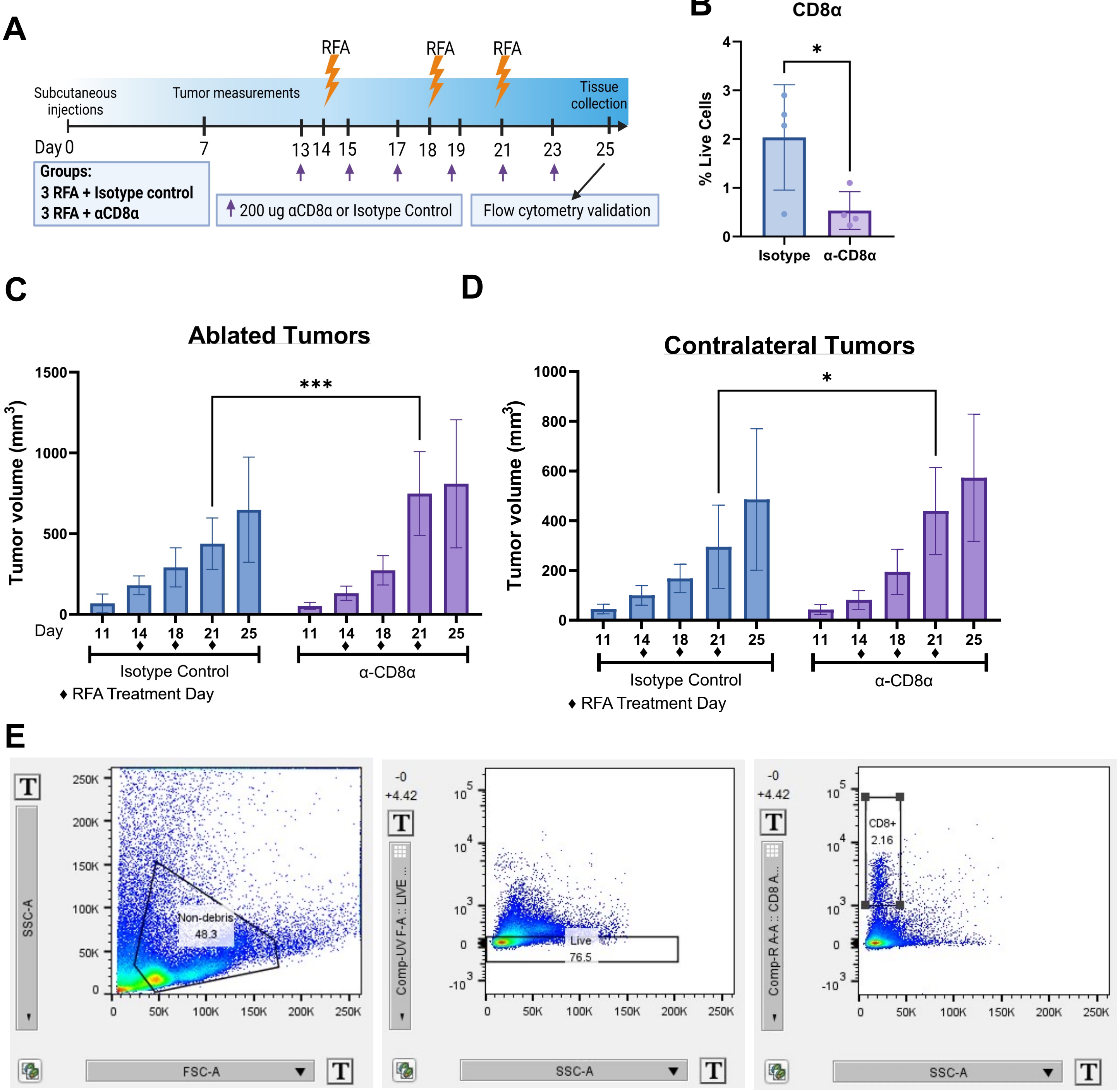


**Supplemental Figure S4. CD8⁺ T cells mediate early tumor control following repeated RFA. A)** Experimental design created using BioRender. **B)** CD8α depletion significantly reduces the %CD8+ cells per live cells in serum. **C)** α-CD8α significantly increases the tumor volume of RFA-treated (***p<0.001) and **D)** contralateral tumors (*p<0.05) after two RFA treatments but does not significantly increase the final tumor volumes after the third RFA treatment. Statistics performed with Prism GraphPad software. n=15 per group. **E)** The data was gated to avoid debris and dead cells while capturing CD8 positive cells. Each sample followed this gating scheme.

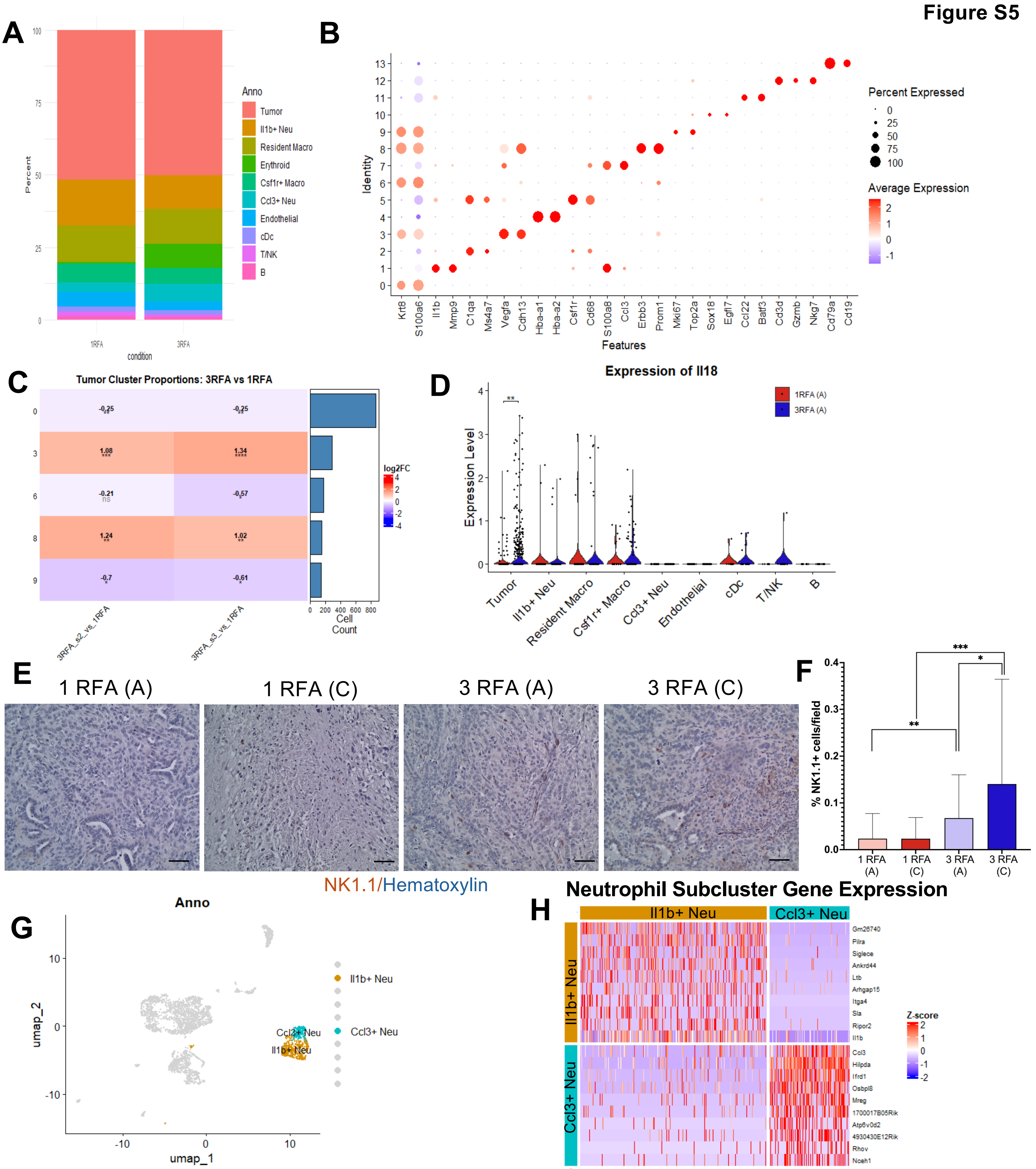


**Supplemental Figure S5. Serial RFA increases natural killer cell infiltration and defines two neutrophil populations. A)** Distribution of cell types as defined by scRNA-seq. **B)** Dot plot of gene expression for each individual cell cluster. **C)** Tumor cluster proportions in 3 RFA and 1 RFA tumors. Clusters 0, 6, 9 were decreased after serial ablation and clusters 3 and 8 were increased after serial ablation. **D)** Violin plot of gene expression of Il18. Il18 levels in the tumor cluster are significantly increased after serial ablations. **E)** Representative 20x images of NK1.1 staining. Scale bars 50 µm. **F)** NK1.1 is significantly increased in both 3 RFA ablated (**p=0.0037) and contralateral (***p=0.0004) tumors compared to respective 1 RFA tumors. NK1.1 staining is significantly increased in 3 RFA contralateral tumors compared to 3 RFA ablated tumors (*p=0.0218). **G)** uMAP of 2 neutrophil clusters. **H)** Heatmap of differentially expressed genes in Il1b+ and Ccl3+ neutrophil clusters.

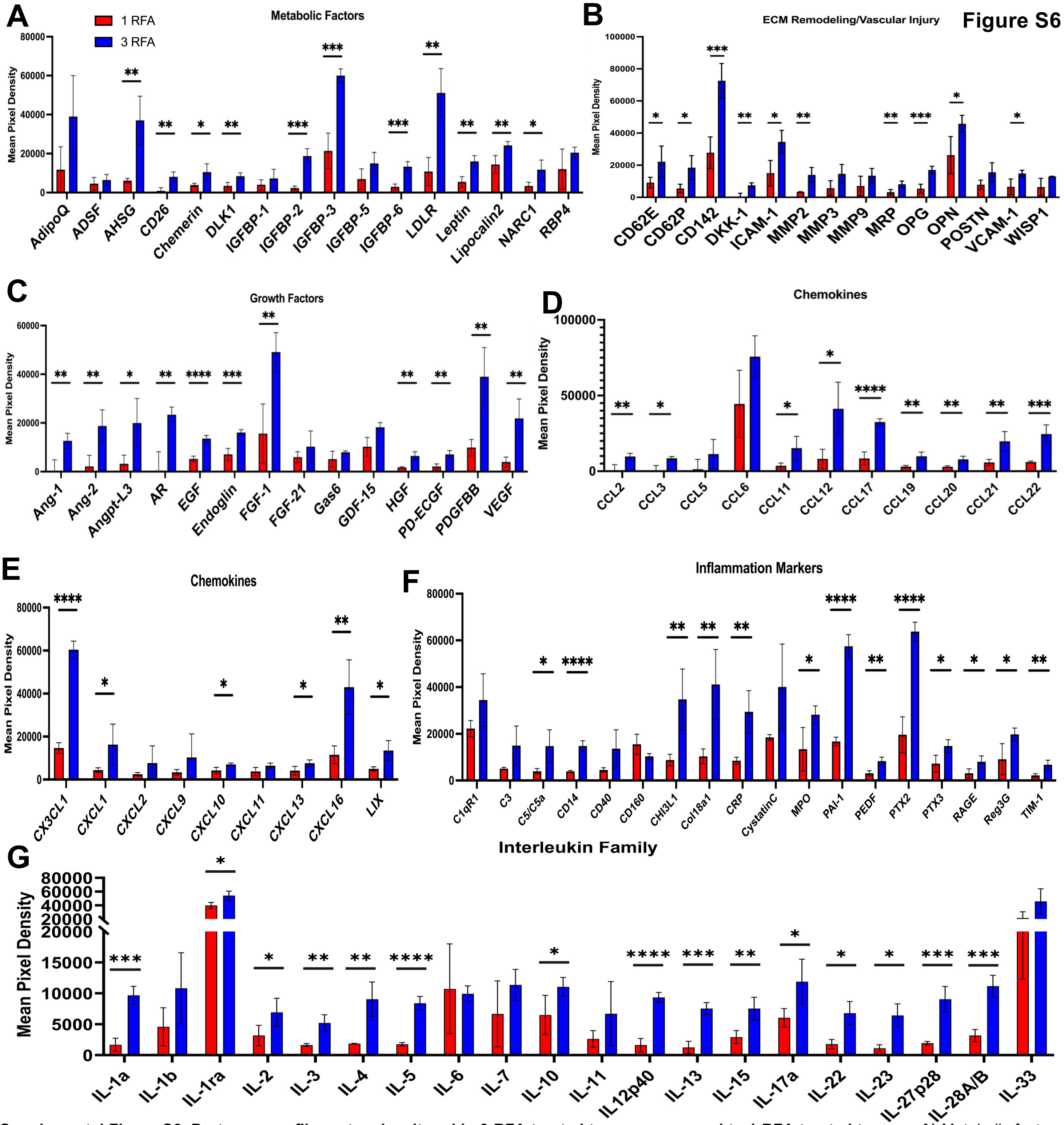


**Supplemental Figure S6. Proteome profiler networks altered in 3 RFA treated tumors compared to 1 RFA treated tumors. A)** Metabolic factors are significantly increased in 3 RFA treated tumors: AHSG (**p=0.0025), CD26 (**p=0.0033), Chemerin (*p=0.0239), DLK1 (**p=0.0063), IGFBP-2 (***p=0.0001), IGFBP-3 (***p=0.0002), IGFBP-6 (***p=0.0005), LDLR (**p=0.0014), Leptin (**p=0.0021), Lipocalin-2 (**p=0.0076), and NARC1 (**p=0.0205). **B)** ECM remodeling/vascular injury proteins are significantly increased in 3 RFA treated tumors: CD62E (*p=0.045), CD62P (*p=0.0176), CD142 (***p=0.0009), DKK-1 (**p=0.0016), ICAM-1 (*p=0.0108), MMP2 (**p=0.0040), MRP (**p=0.0089), OPG (***p=0.0007), OPN (*p=0.0215), and VCAM-1 (*p=0.0206). **C)** Growth factors are significantly increased in 3 RFA treated tumors: Ang-1 (**p=0.0058), Ang-2 (**p=0.0063), Angpt-L3 (*p=0.0202), AR (**p=0.0016), EGF (****p<0.0001), Endoglin (***p=0.0006), FGF-1 (**p=0.0037), HGF (**p=0.002), PD-ECGF (**p=0.0022), PDGFBB (**p=0.0034), and VEGF (**p=0.0049). **D)** CCL family chemokines are significantly increased in 3 RFA treated tumors: CCL2 (**p=0.0098), CCL3 (*p=0.0107), CCL11 (*p=0.026), CCL12 (*p=0.0123), CCL17(****p<0.0001), CCL19 (**p=0.0034), CCL20 (**p=0.0052), CCL21 (**p=0.0064), and CCL22 (***p=0.001). **E)** CXCL family chemokines are significantly increased in 3 RFA treated tumors: CX3CL1 (****p<0.0001), CXCL1 (*p=0.0475), CXCL10 (*p=0.0111), CXCL13 (*p=0.0321), CXCL16 (**p=0.0032), and LIX (*p=0.011). **F)** Inflammation factors are significantly increased in 3 RFA treated tumors: C5/C5a (*p=0.0234), CHI3L1 (****p<0.0001), Col18a1 (**p=0.0068), CRP (**p=0.0039), MPO (*p=0.026), PAI-1 (****p<0.0001), PEDF (**p=0.0019), PTX2 (****p<0.0001), PTX3 (*p=0.0145), RAGE (*p=0.0214), Reg3a (*p=0.0247), and TIM-1 (**p=0.0038). **G)** Interleukin Family proteins significantly increased in 3 RFA treated tumors include: IL-1a (***p=0.0001), IL-1ra (*p=0.0112), IL-2 (*p=0.0379), IL-3 (**p=0.0017), IL-4 (**p=0.0021), IL-5 (****p<0.0001), IL-10 (*p=0.0416), IL12p40 (****p<0.0001), IL-13 (***p=0.0001), IL-15 (**p=0.0045), IL-17a (*p=0.0251), IL-22 (*p=0.0027), IL-23 (*p=0.0017), IL-27p28 (***p=0.0005), IL-28A/B (***p=0.0002). A student's t-test in Graphpad Prism was used for statistical comparisons.

| Target | Application | Primary or Secondary | Company | Catalog Number | Dilution |
|---|---|---|---|---|---|
| NIMPR14/GR1 | IHC | Primary | Abcam | Ab2557 | 1:100 |
| iNOS | IHC | Primary | Abcam | ab15323 | 1:100 |
| CD163 | IHC | Primary | Abcam | Ab182422 | 1:500 |
| Csf1r | IHC | Primary | Abcam | Ab254357 | 1:150 |
| NK1.1 | IHC | Primary | Cell Signaling Technology | E6Y9G | 1:50 |
| Anti-rat | IHC | Secondary | Vector Laboratories | BA-9400-1.5 | 1:500 |
| Anti-rabbit | IHC | Secondary | Vector Laboratories | BA-1000-1.5 | 1:500 |
| CD4 | IF | Primary | Abcam | Ab288724 | 1:35 |
| CD8α | IF | Primary | R&D Systems | MAB116-100 | 1:50 |
| GZMB | IF | Primary | R&D Systems | AF1865 | 1:100 |
| FITC | IF | Secondary | ThermoFisher | A16000 | 1:1000 |
| Cy5 | IF | Secondary | ThermoFisher | A10525 | 1:500 |
| Texas Red | IF | Secondary | ThermoFisher | T-2767 | 1:350 |

**Supplemental Table 1. List of antibodies utilized for immunohistochemistry and immunofluorescence.**
